# Closed-head injury produces a delayed, region-specific defect in glucose-carbon routing at the pyruvate branchpoint in APP/PS1 KI mice

**DOI:** 10.64898/2026.09.29.755475

**Authors:** Elika Z. Moallem, Ronald C. Bruntz, MaKayla F. Cox, Leke Bytyqi, Taylor K. Roberts, Samir P. Patel, Teresa Macheda, Kelly N. Roberts, Emma K. Higgins, Ramon C. Sun, Matthew S. Gentry, Patrick G. Sullivan, Lance A. Johnson, Adam D. Bachstetter

## Abstract

Traumatic brain injury (TBI) is a risk factor for Alzheimer’s disease (AD), and both conditions are associated with impaired glucose metabolism and mitochondrial dysfunction. We previously reported time-dependent mitochondrial dysfunction following closed-head injury (CHI) in amyloid precursor protein/presenilin 1 (APP/PS1) knock-in (KI) mice, but whether glucose-derived carbon labeling showed a corresponding temporal pattern remained unknown. We hypothesized that early deficits would involve tricarboxylic acid (TCA)-cycle metabolites, whereas chronic changes persisted upstream at the pyruvate-lactate branchpoint. Here, oral uniformly ^13^C-labeled glucose ([U-^13^C]-glucose) tracing and gas chromatography-mass spectrometry (GC-MS)-based isotopologue analysis were used to quantify first-turn glucose-derived carbon fate in cortex and hippocampus. To distinguish the effects of age at injury from those of post-injury interval, mice were analyzed 1 month after injury either at 7 or 12 months of age or 8 months after injury at 12 months of age. The most pronounced alterations occurred 8 months after CHI, when KI mice exhibited increased pyruvate and lactate labeling in the cortex and broader increases in glycolytic labeling in the hippocampus. These findings suggest that in APP/PS1 KI mice, chronic closed-head injury produced a delayed shift in glucose-derived carbon labeling centered on glycolytic and pyruvate/lactate nodes, with loss of hippocampal precursor-product coupling but without broad reduction of first-turn TCA-linked labeling.

**Graphical abstract:** 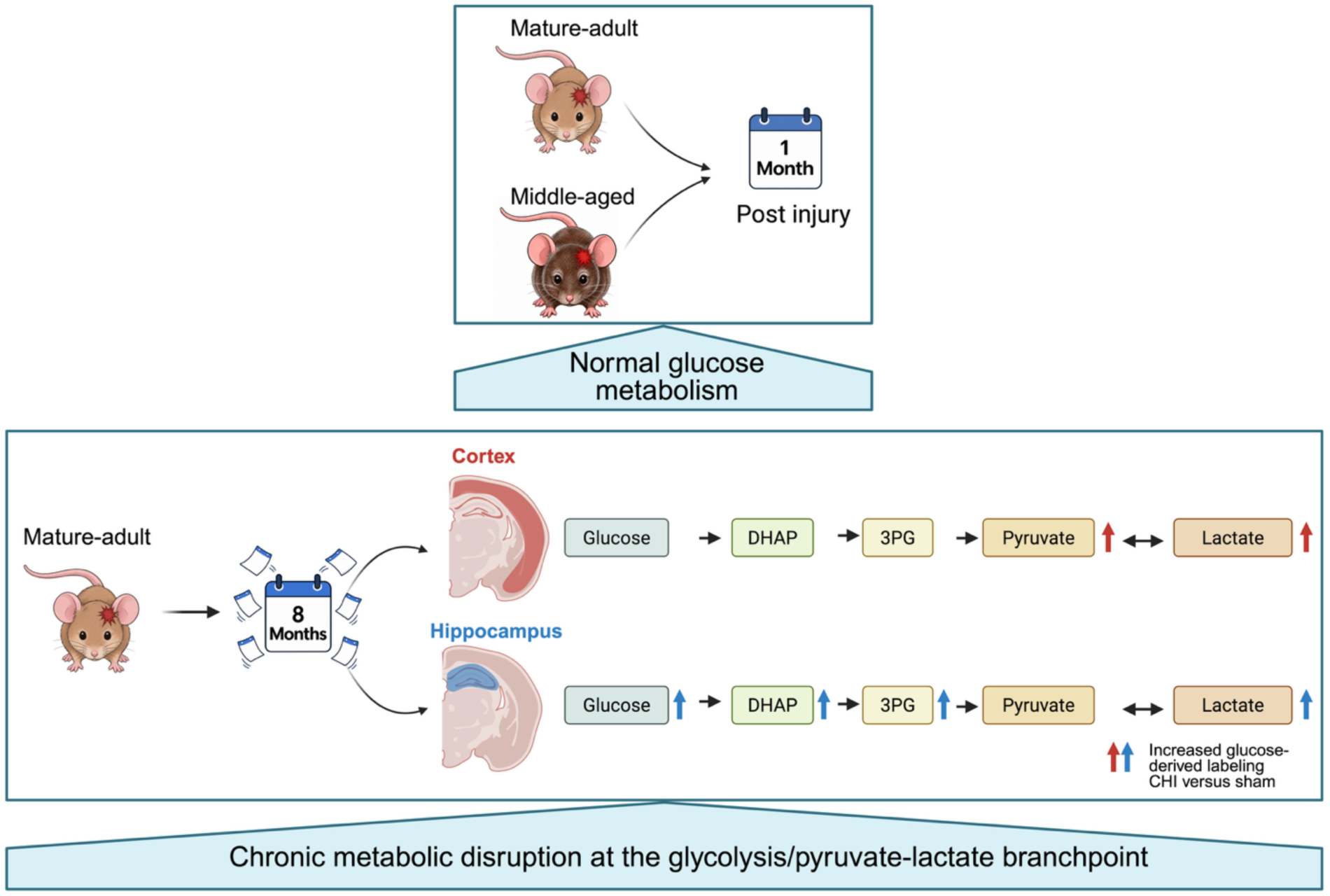

## 1. Introduction

Traumatic brain injury (TBI) is an established risk factor for later Alzheimer’s disease (AD), and both conditions show early deficits in glucose utilization and mitochondrial function (Hagos et al., 2019; Lou et al., 2018; Moallem et al., 2026). Together, these shared deficits suggest that a common metabolic vulnerability may lie in how glucose-derived carbon is handled at the transition from glycolysis to oxidative metabolism. Following injury, cerebral hypometabolism may reflect reduced glucose utilization, impaired downstream routing of glucose-derived carbon after glycolytic entry, or both. How these processes unfold may depend on age at injury and amyloid pathology because age may affect the brain’s capacity to tolerate and recover from metabolic stress, whereas amyloid pathology may further disrupt glucose-carbon metabolism and exacerbate injury-induced deficits. Consequently, how glucose-derived carbon is distributed across downstream metabolic nodes following TBI in an AD-vulnerable brain remains unclear.

Isotope-tracing studies in amyloid-bearing AD mouse models have shown that these defects are node-specific rather than a uniform loss of glucose entry (Sancheti, Kanamori, et al., 2014; Zhou et al., 2018). Labeling of TCA-cycle, neurotransmitter, and pyruvate-linked metabolites changes while glucose carbon continues to enter glycolysis. Several TBI studies describe similar metabolic changes, including lactate accumulation, altered lactate/pyruvate balance, and dysregulated pyruvate dehydrogenase (PDH) across injury models and severities (Carpenter et al., 2015; Lazzarino et al., 2019; Xing et al., 2009). PDH is particularly relevant to this shared vulnerability because it links glycolysis to oxidative metabolism by converting pyruvate to acetyl-CoA, thereby controlling the entry of glucose-derived carbon into the TCA cycle. Tracer studies in acute TBI have further shown increased glucose accumulation alongside reduced downstream oxidative labeling and shunting of carbon into alternative pathways such as the pentose phosphate pathway (Bartnik, Lee, et al., 2007). In the injured human brain, focal 1,2-^13^C_2_ glucose supplementation elevated extracellular pyruvate and lactate without producing detectable TCA-cycle-derived glutamine enrichment, indicating that increased glycolytic flux failed to translate into oxidative carbon entry (Stovell et al., 2023). These effects may be further modified by age at injury. The aged brain shows progressive declines in mitochondrial capacity and metabolic flexibility that are absent or minimal in the mature adult brain, allowing the same injury to produce different metabolic outcomes depending on when it occurs and how long the brain must respond to the resulting routing deficit (Mattson & Arumugam, 2018; Pandya et al., 2015; Yonutas et al., 2015). Although prior ^13^C-tracing studies have examined brain metabolism following TBI and other neurological models, pathway-resolved [U-^13^C]-glucose tracing has not been reported in a closed-head mouse TBI model with an APP/PS1 knock-in (KI) genotype (Bartnik, Lee, et al., 2007; Bartnik et al., 2005; Bartnik-Olson et al., 2010; Clausen et al., 2011; Dusick et al., 2007; Gallagher et al., 2009; Scafidi et al., 2009; Shijo et al., 2017). As a result, it remains unclear whether injury in KI mice produces a distinct glucose-derived carbon-labeling phenotype beyond that seen with TBI alone.

Stable isotope-resolved metabolomics can help address this question because, unlike bulk metabolite measurements, isotopologue patterns reveal whether glucose-derived labeling is preserved, redistributed, or lost across connected metabolic nodes. Oral [U-^13^C]-glucose gavage provides robust labeling of central carbon metabolites in mouse brain and enables analysis of glycolytic, TCA-cycle, and neurotransmitter-linked intermediates over a short first-turn window (Williams et al., 2020). First-turn isotopologue structure has proven informative across disease settings, including subtype-specific routing differences in pediatric tumors (Johnston et al., 2021). Therefore, we applied this rationale in our TBI and AD-relevant model to move beyond the broad label of cerebral hypometabolism and localize where carbon handling becomes impaired.

We recently reported that mature adult KI mice subjected to closed-head injury (CHI) develop time-dependent mitochondrial dysfunction, with the strongest genotype-by-injury effects detected at 1 month post-injury, particularly in the hippocampus (Moallem et al., 2026). Whether this bioenergetic phenotype was accompanied by altered downstream distribution of glucose-derived carbon remained unresolved. Here, we used oral [U-^13^C]-glucose tracing to quantify first-turn isotopologue labeling in the cortex and hippocampus of wild-type (WT) and KI mice subjected to sham or CHI. To distinguish the contributions of age at injury, post-injury interval, and amyloid background, we examined three cohorts: mice injured at 6 months and assessed 1 month later at 7 months of age; mice injured at 11 months and assessed 1 month later at 12 months of age; and mice injured at 4 months and assessed 8 months later at 12 months of age. The first two cohorts allowed comparison of 1-month post-injury responses after injury during mature adulthood versus middle age, whereas the latter two allowed comparison of 1- and 8-month post-injury outcomes at a common 12-month endpoint age. Across all three groups, comparisons between WT and KI mice tested whether amyloid background modified the metabolic response to injury. In KI mice, that amyloid background is accompanied by a local glial response, with microglia and astrocytes accumulating around plaques, where microglia form a barrier that limits protofibrillar Aβ42 and promotes plaque compaction (Condello et al., 2015; Yuan et al., 2016). Because this glial response could itself alter local metabolism, we also quantified amyloid burden and glial marker burden in whole-region and plaque-associated compartments in the same animals.

Based on our previous observation of mitochondrial dysfunction after CHI and the established role of PDH in linking glycolysis to mitochondrial oxidative metabolism, we hypothesized that this subacute mitochondrial phenotype would be accompanied by reduced first-turn glucose-derived labeling of TCA-cycle metabolites relative to upstream glycolytic labeling (Moallem et al., 2026). We predicted that these effects would be more pronounced in mice injured in middle age and in APP/PS1 KI mice. In contrast, because mitochondrial function was no longer altered at 8 months post-injury in mice injured during mature adulthood in our prior study, we did not predict broad reduction in TCA-cycle labeling at that time point. Instead, we predicted that any chronic alteration in glucose-derived carbon handling would persist upstream of oxidative metabolism, particularly at pyruvate- and lactate-linked nodes.

## 2. Methods

### 2.1 Animals

All experimental procedures involving animals were approved by the University of Kentucky Institutional Animal Care and Use Committee and were conducted in accordance with the Guide for the Care and Use of Laboratory Animals and ARRIVE guidelines. APP^NLh/NLh^ × PS1^P264L/P264L^ KI mice (RRID:MGI:3611210), originally developed by Cephalon (Flood et al., 2002), were used in this study.

This strain was generated using a gene-targeted knock-in strategy to introduce the Swedish FAD K670N/M671L mutations, humanize the mouse beta-amyloid sequence (NLh), and insert the P264L mutation into the mouse PS1 gene (Reaume et al., 1996; Siman et al., 2000). Mice were maintained on a CD-1/129 genetic background (RRID:IMSR_CRL:022 × RRID:IMSR_JAX:002448). WT controls were derived from heterozygous APP/PS1 breeding pairs and maintained as a separate lineage, with WT and KI lines backcrossed on average every five generations. Only male mice were included in this study because of animal and tissue availability at the time the cohorts were generated. Prior work from our group suggests that WT mice on the CD-1/129 background are relatively more resistant to TBI-induced mitochondrial changes than WT C57BL/6J mice (Lyons et al., 2018; Moallem et al., 2026). Mice were group housed under a 12 h light/12 h dark cycle with ad libitum access to standard chow and water. Animals were assigned to four experimental groups (WT sham, WT CHI, KI sham, and KI CHI) and studied in three cohorts defined by post-injury interval and age at tissue collection: (1) acute-mature, analyzed 1 month post-injury at approximately 7 months of age; (2) acute middle-aged, analyzed 1 month post-injury at approximately 12 months of age; and (3) chronic middle-aged, analyzed 8 months post-injury at approximately 12 months of age. Mean ages (± SD) at surgery were 5.84 ± 0.12 months for acute-mature, 11.23 ± 0.63 months for acute-aged, and 4.30 ± 0.09 months for chronic-aged. Mean ages at tissue collection were 6.81 ± 0.11 months, 12.19 ± 0.62 months, and 12.30 ± 0.85 months, respectively. Because this was an exploratory, hypothesis-generating study and preliminary isotopologue-tracing data were not available in this model to support a formal a priori power calculation, sample sizes were determined based on cohort feasibility and sample availability. This range is consistent with group sizes commonly used in related mouse injury and metabolomics studies, and we expected that n = 6 per group would be adequate to detect biologically meaningful metabolic differences of moderate-to-large magnitude. Final group sizes ranged from 6 to 11 mice per group.

### 2.2 Closed head injury

Closed head injury (CHI) was induced as previously described (Bachstetter et al., 2015; Lyons et al., 2018; Macheda et al., 2022; Webster et al., 2015). Mice from the different genotype and surgery groups were represented within each surgical session rather than handled in pure group-specific blocks, and the surgeon remained blinded to genotype throughout the procedure. The three time-point cohorts were generated in separate experimental blocks. Within each cohort, mice from all four groups (WT sham, WT CHI, KI sham, and KI CHI) were included in each surgical session, and group assignment was balanced across sessions. The surgeon remained blinded to genotype throughout the procedure. Mice were anesthetized with isoflurane (2.5–4.0%) and secured in a stereotaxic frame. After a midline sagittal incision, a 1 mL water-filled latex pipette bulb (Thermo Fisher Scientific, Waltham, MA, USA) was placed beneath the head. A stereotaxic electromagnetic impactor fitted with a 5.0 mm flat steel tip (Leica Biosystems, Wetzlar, Germany) was then used to deliver a single midline impact to the intact skull at ML 0.0 mm and AP −1.6 mm, with an impact depth of 1.2 mm, velocity of 5.0 ± 0.2 m/s, and dwell time of 100 ms. Sham mice underwent the same anesthesia and incision procedures without impact. Animals were monitored and the righting reflex was recorded during recovery and returned to group housing once full ambulation resumed.

### 2.3 Gavage of [U-^13^C] Glucose solution

^13^C glucose (Cambridge Isotope Laboratories, Tewksbury, MA, USA) was dissolved in ddH2O (Millipore Milli-Q, Bedford, MA, USA) based on the average mouse cohort bodyweight (2g [U-^13^C]-glucose/kg bodyweight). Mice were fasted for 2 h then administered a 2g/kg glucose solution via oral gavage. Brains were collected 30 min after oral gavage of ^13^C glucose. The delivery method and optimal isotopic enrichment time post-gavage was previously optimized (Williams et al., 2020). Throughout this study, “first-turn” labeling refers to the pattern of ^13^C enrichment expected in glycolytic intermediates, pyruvate-branchpoint metabolites, and downstream TCA-linked pools after a single metabolic passage of uniformly labeled glucose carbon, as captured at the 30 min post-gavage time point.

### 2.4 Tissue Collection

Following 30-minute post-gavage interval, mice were euthanized by cervical dislocation and decapitation, and brains were rapidly removed. Brains were bisected sagitally and the hemisphere designated for immunofluorescence was drop-fixed overnight in 4% paraformaldehyde, cryoprotected in 30% sucrose, and sectioned coronally at 30 μm using a freezing microtome. Sections were stored in cryoprotectant at -20°C until staining. Approximately every tenth section between 1.3mm and 2.5mm posterior to bregma was collected for staining. Two sections per animal were stained and quantified for each marker. Cortex tissue designated for biochemical assays was frozen, pulverized, and stored at -80°C until extraction. Samples designated for GC-MS, multiplex immunofluorescence staining, and MSD were randomized before assay-specific preparation.

### 2.5 Multiplex Immunofluorescence Staining

To assess the spatial relationship between IBA1-positive microglia, CD45-positive signal, and plaques, multiplex immunofluorescence was performed on free-floating coronal sections from the same animals using systematic sampling. Sections were sampled at approximately every one-in-ten intervals between 1.3 mm and 2.5 mm posterior to bregma, and two 30 μm sections per animal were stained and quantified for each marker. One section included both neocortex and hippocampus and the second section only neocortex. Slides were scanned using a Zeiss AxioScan Z.1.

Free-floating sections were washed three times in 1X PBS for 10 minutes at room temperature with shaking and blocked for 60 minutes at room temperature in goat blocking buffer containing 10% goat serum, 0.2% Triton X-100, and 1X PBS. Sections were incubated for 24 hours at 4°C with gentle shaking in primary antibodies against CD45 (rat anti-CD45, BioLegend 103102, RRID:AB_312967, 1:200), IBA1 (guinea pig anti-IBA1, HistoSure Synaptic Systems HS-234308, RRID:AB_2924932, 1:1000), and GFAP (rabbit anti-GFAP, Dako Z0334, RRID: AB_10013382, 1:1,000). The antibody diluent contained 3% goat serum, 0.2% Triton X-100, and 1X PBS.

Sections were washed five times in 1X TBS-T for 5 minutes and incubated for 1 hour at room temperature with fluorophore-conjugated secondary antibodies protected from light: Alexa Fluor 555 goat anti-rat IgG (1:500), Alexa Fluor 488 goat anti-guinea pig IgG (1:200), and Alexa Fluor 647 goat anti-rabbit IgG (1:500). Sections were then washed three times in 1x PBS for 10 minutes, incubated for 5 minutes in 1X Trueblack lipofuscin quencher (Biotium 23007), mounted onto slides, and air-dried in the dark for 15 minutes. Sections were then incubated with 1x Amylo-Glo for 10 minutes and coverslipped using Invitrogen Diamond Antifade Mounting Medium.

### 2.6 Plaque associated analysis

Plaque-association analysis was performed on the multiplex immunofluorescence images to quantify Aβ plaque-associated and non-plaque-associated marker-positive areas using HALO Area Quantification FL v3.0.1. Whole-region annotations were manually defined for neocortex and hippocampus. Amylo-Glo-positive plaque cores were identified using the DenseNet V2 HALO AI classifier. As previously applied in this laboratory (Shahidehpour et al., 2023), concentric partitioners were generated outward from the plaque boundaries to define a 0-10 μm annulus, with an exclusion annotation used to remove the plaque core from the 0-10 μm annulus. An additional negative concentric partitioner defined tissue more than 50 μm from the plaque as the non-plaque-associated reference compartment. All annotations were visually reviewed before quantification.

Images were analyzed in HALO (Indica Labs) using fixed analysis settings across slides. Area quantification was performed for the Amylo-Glo, CD45, IBA1, and GFAP channels. Plaque-compartment analyses were retained separately from whole-region analyses.

### 2.7 Sample preparation for GC-MS analysis

#### Tissue Pulverization and Extraction

Mice were euthanized by cervical dislocation followed by decapitation. Brains were rapidly removed and dissected into the neocortex and hippocampus, frozen in liquid nitrogen, and stored at −80°C until further processing. Decapitation-to-freezing time was less than 90 seconds for all animals. Collection order was randomized with respect to genotype and surgery group, and samples were processed in batches containing all experimental groups. Metabolites were extracted for mass spectrometry analysis using techniques previously described (Sun et al., 2021; Williams et al., 2020). Frozen cortex and hippocampus tissue were pulverized under liquid nitrogen using a Freezer/Mill Cryogenic Grinder (SPEX SamplePrep, Metuchen, NJ, USA), and the resulting tissue powder was maintained on liquid nitrogen during handling.

Approximately 20 mg of frozen tissue powder was transferred with a pre-chilled micro spatula into a pre-labeled 1.5 mL microcentrifuge tube. An equal volume of 50% methanol containing 20 μM L-norvaline, prepared by adding 1 μL of 20 mM L-norvaline to 1 mL of 50% methanol, was added to each sample. L-Norvaline was included as an internal standard according to the established extraction protocol but was not used to normalize fractional isotopologue enrichment, which was calculated from the isotopic distribution within each metabolite.

Samples were briefly vortexed and placed on ice. Mechanical disruption was performed using a cell disruptor at 3,000 rpm for 3 minutes and incubated on ice for 30 minutes with intermittent vortexing every 10 minutes. Following incubation, samples were centrifuged at 18,000 × g for 10 minutes at 4°C. The supernatant containing the polar metabolite fraction was transferred to a separate tube and dried at 10^-3^ mbar using a SpeedVac concentrator (Thermo Fisher Scientific, Waltham, MA, USA).

All metabolite data reported in this study were obtained from the polar supernatant and therefore represent the free metabolite pool. Any measurements obtained from the hydrolyzed insoluble biomass fraction, including protein-bound amino acids and thymine, were not included in the analyses reported here.

#### Derivatization for GC-MS

Before derivatization, dried polar extracts were returned to the SpeedVac for 20 min to ensure complete removal of residual moisture. Methoxyamine hydrochloride (20mg/mL) was prepared in pyridine, and 50 μL of this solution was added to each dried sample. Samples were vortexed and incubated at 30°C for 90 min. Following incubation, samples were centrifuged at 18,000 x g for 10 min and 40 μL of the supernatant was transferred to an amber glass chromatography vial (Agilent Technologies). N-methyl-N-trimethylsilyl-trifluoroacetamide (MSTFA; Thermo Fisher Scientific) was added to each sample at a volume of 80 μL using a glass pipette. Samples were briefly vortexed and incubated at 37°C for 30 min on a heat block.

##### GC-MS isotopologue analysis

Derivatized samples were analyzed using an Agilent 7890B gas chromatography (GC) instrument coupled to an Agilent 7010A triple-quadrupole mass spectrometer equipped with a high-efficiency source. A 2-μL aliquot of each sample was injected into an HP-5MS UI column (30 m × 250 μm internal diameter, 0.25 μm film thickness) using a 1:5 split ratio. Helium was used as the carrier gas at a constant flow rate of 1.5 mL/min. The mass spectrometer interface and ion-source temperatures were maintained at 290°C and 230°C, respectively, and electron ionization (EI) performed at 70 eV. Chromatographic separation was achieved using a modified thermal gradient. The initial oven temperature was maintained at 60°C for 1 min, increased at 10°C/min to 325°C, then maintained at 325°C for 10 min. Data were acquired in full-scan mode over an m/z range of 50–800. Metabolites were identified by matching their retention times and EI fragmentation patterns to reference spectra in the Fiehn GC-MS Metabolomics Library (Agilent Technologies).

Raw GC-MS data were processed using Data Extraction for Stable Isotope-labelled metabolites (DExSI; (Dagley & McConville, 2018). For each metabolite, peak areas were integrated for the m+0 through m+n ion series of a selected EI fragment containing a defined number of carbon atoms derived from the native metabolite. Because electron ionization produces metabolite-specific fragments, the measured labeling distribution represents the carbon atoms retained within the selected fragment rather than necessarily representing the entire native metabolite. For each metabolite, the selected EI fragment, retention time, monitored isotopologue m/z values, and fragment formula used in IsoCorrectoR for natural-abundance correction are provided in Supplemental Table 1.

Glucose was the one exception and was quantified from two monitored derivative fragments, a 160 m/z fragment containing carbons 1–2 and a 319 m/z fragment containing carbons 3–6, using the fragment assignments described in (Kohlstedt & Wittmann, 2019). Labeling did not differ between the two fragments, consistent with uniform labeling of the glucose molecule. This uniformly labeled measurement is referred to as glucose m+6 throughout.

Exported peak areas were corrected for natural abundance using IsoCorrectoR with default settings (Heinrich et al., 2018). At the nominal mass resolution used in this study, naturally occurring heavy-isotope contributions occurring at the same nominal m/z as tracer-derived mass shifts were not resolved separately. These contributions were therefore addressed through natural-abundance correction of the monitored derivatized fragment. Following correction, the fractional enrichment of each isotopologue was calculated by dividing its corrected peak area by the sum of the corrected peak areas for all measured isotopologues of that fragment.

The resulting values represent corrected mass isotopologue distributions rather than positional isotopomer abundances or absolute metabolite concentrations. Because fractional enrichment reflects the isotopic distribution within each monitored metabolite fragment, the reported values were not normalized to L-norvaline, thymine, or protein content.

### 2.8 Aβ42 quantification by Meso Scale Discovery Assay

Aβ42 was measured in residual pulverized cortex remaining after the GC-MS extraction. Aβ42 was quantified in PBS-soluble and formic-acid-soluble cortical fractions using a singleplex V-PLEX Aβ42 Peptide assay with the 4G8 configuration (Meso Scale Discovery). The assay was performed on an Aβ42 Peptide 96-well 4-spot sector plate (catalog no. N451LBA-1; lot Z0057639; barcode 25P0BATG12; CoA lot K0082850).

The MSD assay used frozen cortical powder that was pulverized prior for the GC-MS workflow. To generate the PBS-soluble fraction, PBS lysis buffer was added at 20x the tissue weight /volume (20 μL/mg tissue), followed by homogenization and centrifugation at 20,000 x g for 20 minutes. The PBS supernatant was collected and analyzed neat. The formic-acid-soluble fraction was generated using 70% formic acid followed by homogenization, centrifugation at 20,000 x g for 30 minutes at 4°C, and collection of the supernatant for storage at -80°C. Formic-acid samples were neutralized in a 1:20 step and then diluted an additional 1:10 in PBS, yielding a cumulative 1:200 dilution immediately before assay. The plate was washed with PBS containing 0.05% Tween-20, then incubated with samples overnight at 4°C with continuous shaking, washed, incubated with detection antibody for 2 hours at room temperature with shaking, washed again, and developed with MSD read buffer before reading. Protein concentration was determined in the PBS fraction using the Pierce BCA Protein Assay Kit (catalog no. 23225). Samples were diluted 1:6, measured in triplicate, combined with BCA reagents A and B at a 50:1 ratio, incubated at 37°C for 30 minutes, and read at 562nm. Both Aβ42 fractions were normalized to the PBS-fraction BCA protein concentration and reported as pg Aβ42/μg protein.

### 2.9 Statistical analysis

Statistical analyses were performed in R version 4.5.2. Data are presented as mean ± SEM unless otherwise stated, and n denotes biologically independent mice. Righting-reflex latency group differences were assessed by one-way ANOVA followed by Tukey post hoc testing. Secondary models evaluated genotype, surgery, and genotype × surgery effects within each cohort. For first-turn isotopologue labeling, overall effects across the metabolite panel were tested by multivariate analysis of variance (MANOVA; Pillai trace) with surgery, genotype, brain region, and cohort as factors. Individual isotopologues were then analyzed by Type III ANOVA using the same factorial structure. Within-stratum CHI-versus-sham comparisons were performed using two-sided Welch two-sample t-tests. For precursor-product coupling analyses, Pearson correlation coefficients were calculated separately for sham and CHI animals within each brain region × genotype × cohort panel. Sham-versus-CHI differences in correlation strength were assessed using Fisher’s r-to-z comparisons. Differences in precursor-product slopes were tested using linear models followed by Type III ANOVA.

Analyses were designated as primary or hypothesis-generating before statistical testing. Primary analyses were the first-turn MANOVA, the follow-up Type III ANOVAs for individual isotopologues, and the planned CHI-versus-sham Welch contrasts within each genotype and brain region. The principal comparison was APP/PS1 KI CHI versus KI sham in the chronic 8-month post-injury cohort. The precursor-product correlation and slope analyses (Figs. 3–5 and S1-S5) were hypothesis-generating. Benjamini-Hochberg correction was applied as a sensitivity analysis rather than as a decision rule. Because family size is a judgement rather than a property of the data, each p-value was corrected under both a narrow and a broad family. Only the ranking set differed, and no model was refit. First-turn labeling p-values were ranked within the chronic cohort’s 48 contrasts and again within all 144 contrasts across the three cohorts, which those 48 are a subset. Family definitions for each analysis are given in Supplemental Table 2.

Immunofluorescence were analyzed by one-way ANOVA followed by pairwise comparisons between KI sham and KI CHI using Mann-Whitney U test. All statistical tests were two-sided, and p < 0.05 was considered statistically significant. Full statistical reporting for all analyses included in the submitted manuscript is provided in Supplemental Table 2.

## 3. Results

### 3.1 Acute neurological impairment after CHI

Righting reflex latency was assessed immediately after CHI as an index of acute neurological impairment and injury severity. Time to regain a prone posture was recorded in seconds from impact until each mouse successfully righted itself. This matched analysis included 78 mice with numeric righting-reflex values from the isotopologue cohort. One-way ANOVA demonstrated a group effect on righting time (F(9, 68)=6.38, p<0.001). Tukey-adjusted post hoc comparisons showed prolonged righting reflex latency in WT CHI mice injured at 11 months (middle-aged, 1-month post-injury; 416.91 s) versus sham controls (90.00 s, Tukey-adjusted p<0.001) and in KI CHI mice injured at 11 months (middle-aged, 1-month post-injury; 410.29 s) versus sham controls (75.83 s, Tukey-adjusted p=0.006).

### 3.2 APP/PS1 KI genotype and post-injury time broadly alter first-turn glucose-carbon fate, with injury specificity for substrate and pyruvate-lactate labeling

To test the hypothesis that CHI alters glucose-derived carbon flow across the first-turn metabolic network, we first used multivariate analysis to determine whether the overall isotopologue profile differed by injury, genotype, post-injury cohort, and brain region. Throughout the Results, the three post-injury cohorts follow the design introduced above (Fig. 1): mature adult, injured at 6 months, 1-month post-injury (7 months at endpoint); middle-aged, injured at 11 months, 1-month post-injury (12 months at endpoint); and mature adult, injured at 4 months, 8-month (chronic) post-injury (12 months at endpoint). We then used univariate analyses to identify which specific metabolites accounted for these network-level effects and to test whether the injury reduced first-turn TCA-cycle labeling relative to upstream glycolytic labeling, as hypothesized. Representative first-turn labeling routes downstream of oral [U-^13^C]-glucose are illustrated in Fig. 2A. At the network level, brain region, genotype, and experimental cohort all significantly influenced first-turn labeling, whereas surgery as a main effect was limited. Multivariate analysis by Pillai’s trace showed strong main effects of brain region (F(12, 132)=33.093, p<0.0001), genotype (F(12, 132)=4.189, p<0.0001), and cohort (F(24, 266)=6.370, p<0.0001), whereas surgery did not exert a significant multivariate effect across the panel (F(12, 132)=1.621, p=0.0931). Univariate Type III ANOVAs further showed broad effects of genotype and cohort across the pathway, with genotype significant for all 12 first-turn isotopologues and cohort significant for 11 of 12 (Fig S1). Region effects were more selective at the individual-metabolite level, involving eight isotopologues spanning glucose m+6, early glycolytic intermediates, alanine m+3, and several downstream mitochondrial-or aspartate-linked nodes, whereas surgery as a main effect was limited to DHAP m+3 (Fig. 2B). Pathway-wide variation was therefore driven primarily by genotype, cohort, and region, with injury effects restricted to selected upstream metabolites. Significant genotype × cohort and region × cohort interactions further indicated that these network-level effects varied across cohorts. By contrast, surgery × cohort effects were limited to glucose m+6 and lactate m+3, suggesting that injury-related differences were cohort dependent and restricted to selected upstream metabolites.

**Figure 1:**
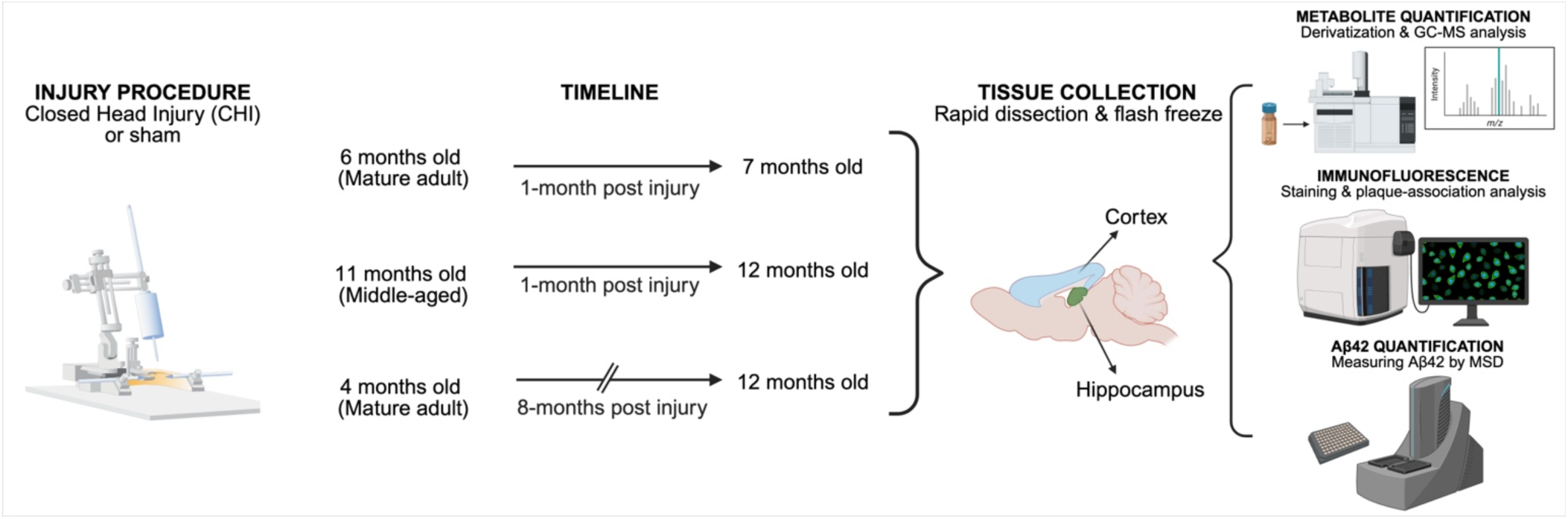
Experimental design in WT and KI male mice across three post-injury cohorts. WT and KI male mice were studied in the mature adult 1-month post-injury cohort (7 months at endpoint), the middle-aged 1-month post-injury cohort (12 months at endpoint), and the chronic 8-months post-injury cohort (mature adult-injured, 12 months at endpoint). Within each cohort, mice were assigned to sham or CHI. Following a 2-hour fast, mice received oral [U-^13^C]-glucose (2 g/kg) and were euthanized 30 minutes later. Cortex and hippocampus were rapidly dissected, flash-frozen, and processed for GC-MS isotopologue analysis. Selected animals were additionally assessed by multiplex immunofluorescence for Aβ plaques, GFAP, IBA1 and CD45, including plaque-association analysis. Selected tissue samples were also analyzed by singleplex MSD for PBS-soluble and formic-acid-soluble AB42. Final post-QC group sizes for the GC-MS dataset were as follows: mature adult 1-month post-injury, WT sham n = 8, WT CHI n = 8, KI sham n = 10, KI CHI n = 8; middle-aged 1-month post-injury, WT sham n = 8, WT CHI n = 11, KI sham n = 6, KI CHI n = 7; chronic 8-month post-injury, WT sham n = 6, WT CHI n = 6, KI sham n = 6, KI CHI n = 6. Multiplex immunofluorescence and singleplex MSD were performed in assay-specific subsets; sample sizes varied by cohort, genotype, surgery, brain region, and MSD fraction because of tissue availability and assay-specific quality-control exclusions. Exact endpoint-specific sample sizes are provided in the corresponding figure legends and Supplemental Table 2.

**Figure 2:**
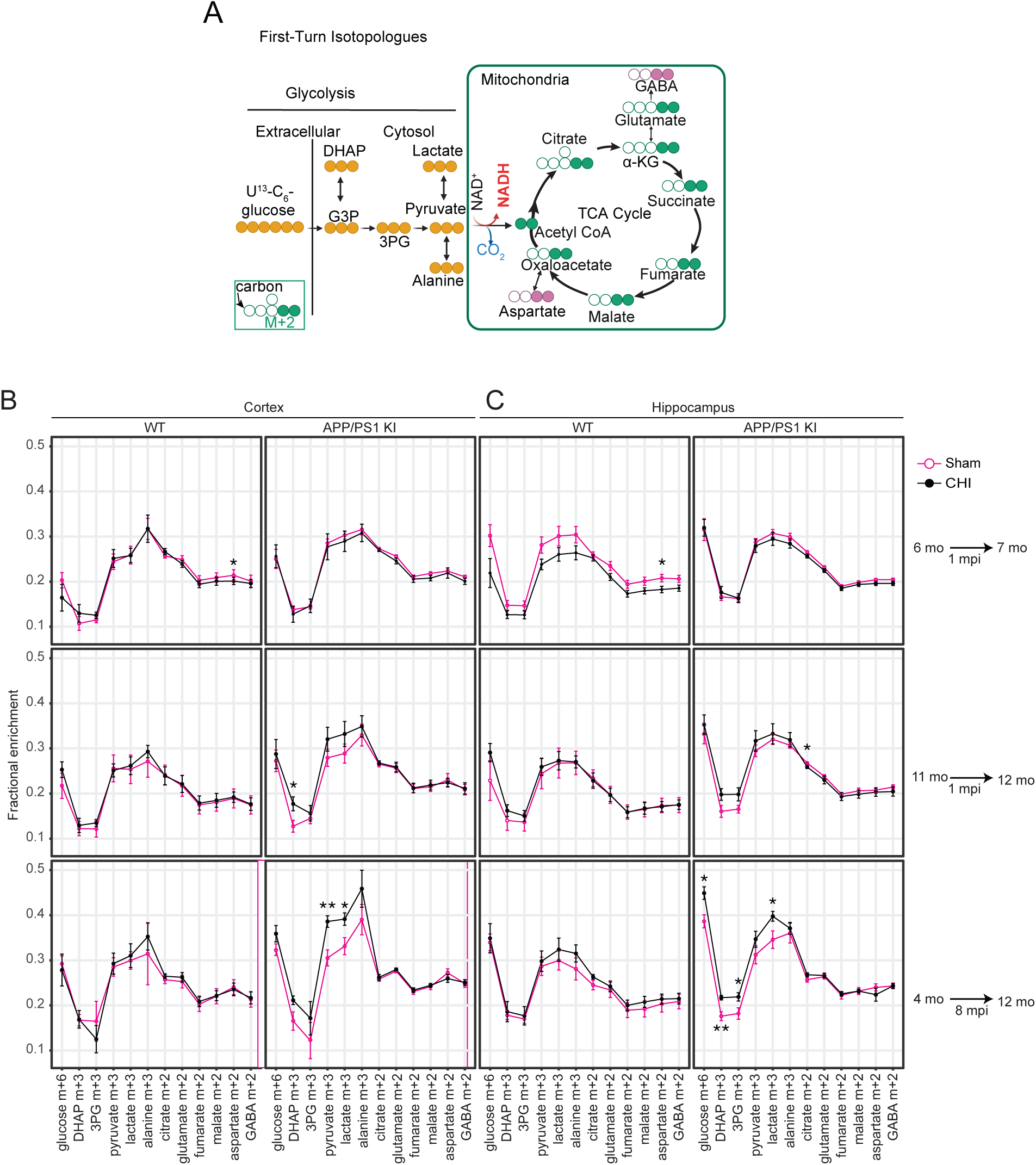
APP/PS1 KI genotype and post-injury cohort broadly reprogram first-turn glucose-carbon fate, with injury effects largely restricted to substrate and pyruvate-lactate labeling. **(A)** Schematic of representative first-turn ^13^C-labeling routes downstream of oral [U-^13^C]-glucose. Label enters through glycolytic intermediates (DHAP, 3PG, pyruvate) and at the pyruvate branchpoint is distributed to lactate via LDH, to alanine via transamination, or to the TCA cycle as acetyl-CoA, generating citrate m+2 and downstream first-turn intermediates including glutamate m+2, fumarate m+2, malate m+2, aspartate m+2, and GABA m+2. Filled circles, ^13^C; open circles, ^12^C. **(B-C)** Mean fractional enrichment (+/-SEM) of first-turn isotopologues plotted in pathway order for cortex **(B)** and hippocampus **(C)**, stratified by genotype (WT, KI) and post-injury cohort (rows: mature adult 1-month post-injury (1 mpi), middle-aged 1-month post-injury (1 mpi), chronic 8-month post-injury (8 mpi)). Asterisks denote two-sided Welch two-sample t-tests comparing CHI versus sham within each genotype x cohort stratum (nominal p values: *p < 0.05, **p < 0.01, ***p < 0.001). n = 6-11 male mice per genotype x surgery x cohort group. Abbreviations: mpi, month post-injury; mo, months.

### 3.3 Chronic, but not acute, CHI effects in APP/PS1 KI mice were restricted to glycolytic and pyruvate/lactate nodes

Because the preceding network-level analysis showed that injury did not exert a broad main effect on first-turn glucose-carbon flow, we next asked whether CHI-related differences emerged only within specific genotype- and cohort-defined comparisons. These within-genotype CHI-versus-sham contrasts directly tested our prediction that acute injury would be associated with deficits in TCA-cycle labeling, particularly in middle-aged and APP/PS1 KI mice, while chronic post-injury abnormalities would appear at upstream glycolytic and pyruvate/lactate-linked nodes. At 1 month post-injury, injury-related differences were limited and did not support the predicted acute pattern of broad TCA-cycle impairment. In the mature-adult cohort injured at 6 months (1-month post-injury), the only significant difference was in WT hippocampus, where CHI reduced aspartate m+2 relative to sham (sham mean=0.208, CHI mean=0.183; t(13.72)=2.315, p=0.0366), an isolated downstream finding without accompanying changes in upstream glycolytic or branchpoint metabolites. In the middle-aged cohort injured at 11 months (1-month post injury), KI cortex showed higher DHAP m+3 in CHI than in sham (sham=0.128, CHI=0.177; t(9.76)=-2.445, p=0.0351), whereas KI hippocampus showed lower citrate m+2 in CHI (sham=0.268, CHI=0.259; t(7.54)=2.569, p=0.0348). Between the two 1-month cohorts, the affected node, brain region and direction of change all differed. No other isotopologues reached nominal significance in either cohort.

In contrast, at 8 months post-injury, CHI-versus-sham differences in KI brain were larger and more internally consistent. This pattern was consistent with the prediction that chronic abnormalities would be expressed upstream of oxidative metabolism rather than as broad TCA-cycle deficits. Every significant KI difference at this time point showed increased labeling, although the affected metabolites differed between cortex and hippocampus. In KI cortex, CHI selectively elevated pyruvate m+3 (sham=0.305, CHI=0.386; t(9.10)=-3.679, p=0.00499) and lactate m+3 (sham=0.331, CHI=0.391; t(9.15)=-2.564, p=0.0301), whereas upstream glycolytic intermediates (glucose m+6, DHAP m+3, 3PG m+3) were not significantly altered. In KI hippocampus, the pattern extended further upstream: CHI elevated glucose m+6 (sham=0.386, CHI=0.449; t(9.99)=-3.115, p=0.0110), DHAP m+3 (sham=0.176, CHI=0.217; t(7.50)=-3.699, p=0.00679), 3PG m+3 (sham=0.182, CHI=0.219; t(9.15)=-2.357, p=0.0423), and lactate m+3 (sham=0.346, CHI=0.398; t(7.83)=-2.320, p=0.0496). Pyruvate m+3 did not significantly differ in KI hippocampus at this time point (sham=0.312, CHI=0.347; p=0.228), indicating that the upstream accumulation detected at glucose, DHAP, and 3PG did not extend as a statistically clear increase to pyruvate itself. By contrast, downstream first-turn TCA-cycle intermediates, including citrate m+2, glutamate m+2, fumarate m+2, malate m+2, aspartate m+2, and GABA m+2, showed no significant CHI-related differences in either KI cortex or KI hippocampus. No significant CHI-related differences were detected in WT cortex or hippocampus at 8 months post-injury. Cortical accumulation localized to the pyruvate/lactate branchpoint, while hippocampal accumulation extended from glucose through 3PG and lactate (Fig. 2B–C; see also, Supplemental Table 2).

### 3.4 CHI-versus-sham coupling differences are localized to the pyruvate branchpoint and did not extend to downstream TCA-linked precursor-product pairs

After identifying chronic KI CHI accumulation at glycolytic and pyruvate/lactate nodes, we next examined correlations between linked metabolite pairs to localize where carbon flow was preserved versus lost. As an exploratory test of carbon labeling through lower glycolysis, we examined the relationship between 3PG m+3 and pyruvate m+3 (Fig. 3A). Across the two 1-month post-injury cohorts, positive coupling was present in most groups, particularly in hippocampus, so coupling from 3PG to pyruvate was generally preserved (Fig. 3B). In the chronic 8-month post-injury cohort, this relationship was largely retained in WT tissue but was lost in KI tissue. In KI cortex, neither sham nor CHI animals showed significant coupling, whereas in KI hippocampus sham animals retained a strong positive correlation but CHI animals showed no detectable association. The strongest injury-associated change in 3PG-to-pyruvate coupling was therefore in chronic KI hippocampus, consistent with disruption of carbon flow at this glycolytic step.

**Figure 3:**
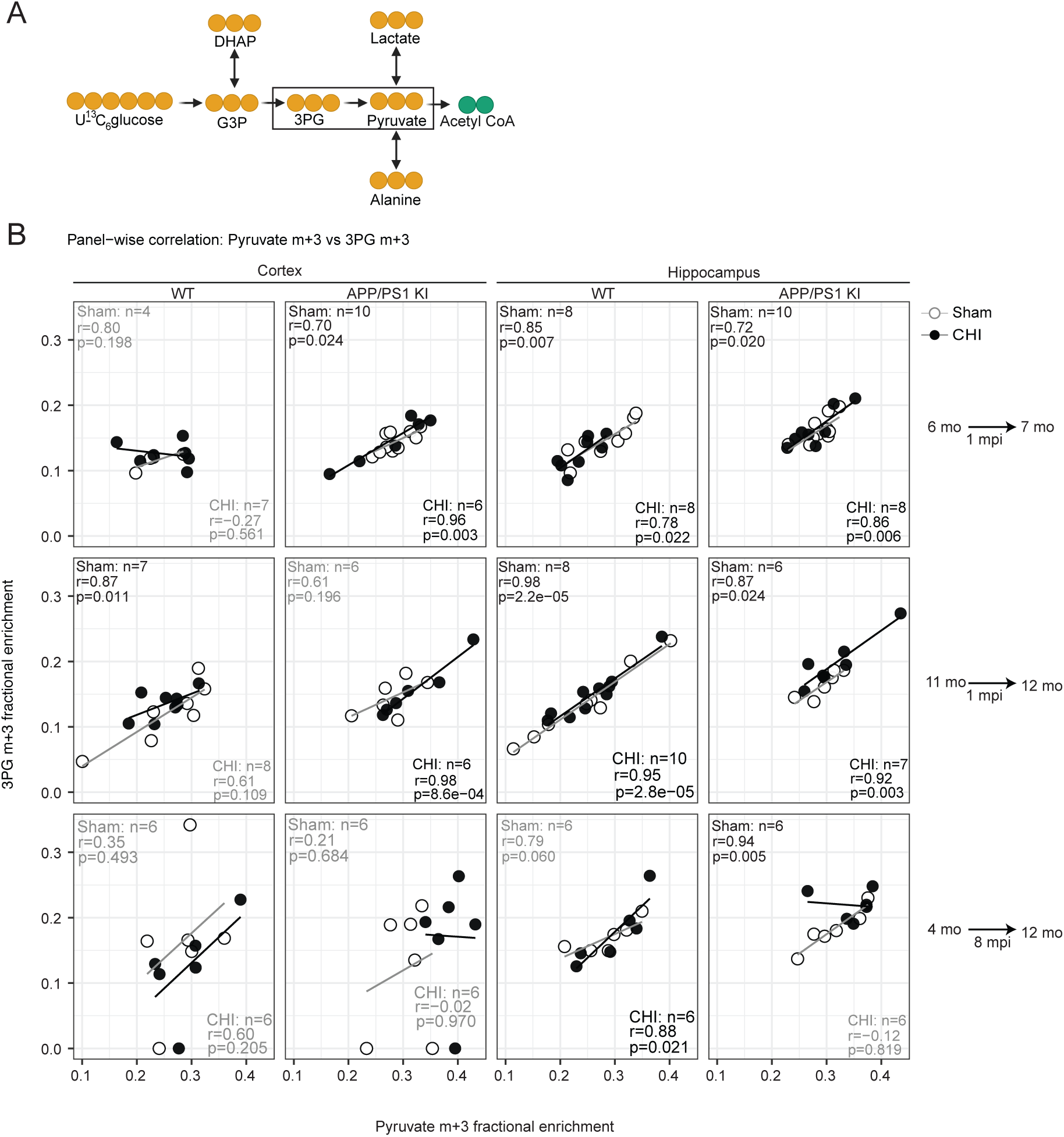
Loss of 3PG-to-pyruvate coupling in chronic KI hippocampus after CHI. (**A**) Schematic showing the 3PG-to-pyruvate step within lower glycolysis and its position relative to pyruvate branchpoint fates. (**B**) Panel-wise correlations between pyruvate m+3 and 3PG m+3 fractional enrichment in cortex and hippocampus from WT and KI mice under sham or CHI conditions across three cohorts: mature-adult 1-month post-injury (7 months at endpoint), middle-aged 1-month post-injury (12 months at endpoint), and chronic 8-month post-injury (mature-adult-injured, 12 months at endpoint). Sham-versus-CHI differences in correlation strength were assessed by Fisher’s r-to-z comparison, and differences in regression slope were assessed using a slope-interaction linear model (y ∼ x * surgery). Abbreviations: mpi, month post-injury; mo, months.

Because pyruvate is a branchpoint between lactate production and oxidative metabolism, we examined pyruvate m+3 and lactate m+3 coupling as an exploratory measure of branchpoint metabolism (Fig. 4A). Across nearly all groups, pyruvate m+3 and lactate m+3 remained strongly and positively correlated, so this lactate dehydrogenase (LDH)-linked relationship was generally preserved (Fig. 4B). The major exception was KI hippocampus in the chronic 8-month post-injury cohort, where sham animals retained strong positive coupling, but CHI animals showed no detectable association. Combined with the loss of 3PG-to-pyruvate coupling in the same group, chronic KI CHI hippocampus showed disrupted carbon metabolism around the pyruvate branchpoint.

**Figure 4:**
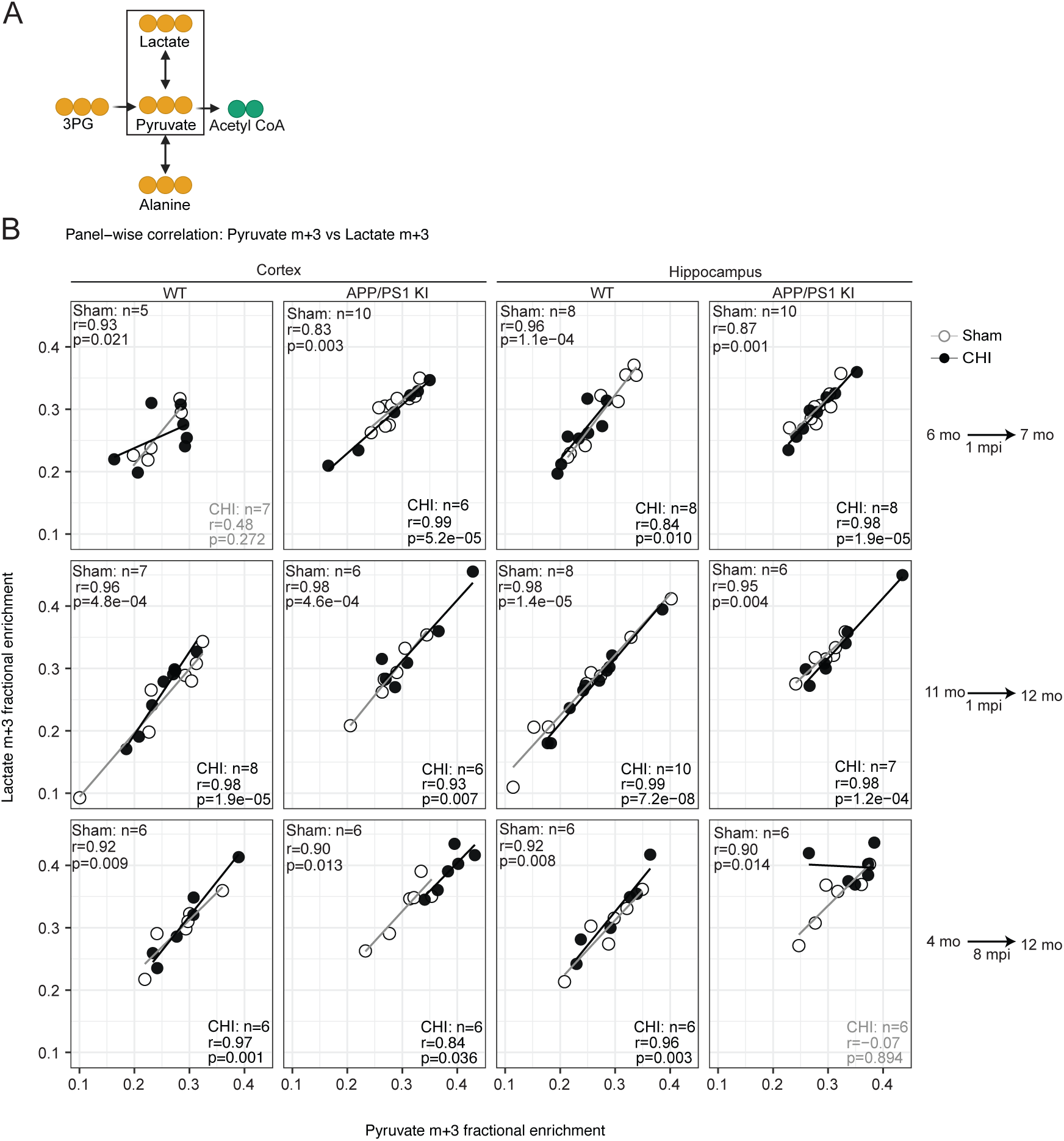
Selective loss of pyruvate-to-lactate coupling in chronic KI hippocampus after CHI. **(A)** Schematic showing the reversible pyruvate-to-lactate branchpoint and its position relative to pyruvate entry into oxidative metabolism. **(B)** Panel-wise correlations between pyruvate m+3 and lactate m+3 fractional enrichment in cortex and hippocampus from WT and KI mice under sham or CHI conditions across three cohorts: mature-adult 1-month post-injury (7 months at endpoint), middle-aged 1-month post-injury (12 months at endpoint), and chronic 8-month post-injury (mature-adult-injured, 12 months at endpoint). Sham-versus-CHI differences in correlation strength were assessed by Fisher’s r-to-z comparison, and differences in regression slope were assessed using a slope-interaction linear model (y ∼ x * surgery). Abbreviations: mpi, month post-injury; mo, months.

Alanine labeling reflects pyruvate transamination rather than reversible pyruvate-lactate exchange through lactate dehydrogenase (Fig. 5A). Pyruvate m+3 and alanine m+3 were positively coupled in only a subset of conditions, so carbon transfer from pyruvate to alanine was less consistently maintained than pyruvate-to-lactate coupling (Fig. 5B). Across the two 1-month post-injury cohorts, positive pyruvate-to-alanine coupling was present in several groups, with WT hippocampus showing the most consistent association. In the chronic 8-month post-injury cohort, this relationship remained positive in WT tissue but was weakened in KI CHI tissue. In both KI cortex and KI hippocampus, sham animals retained positive pyruvate-to-alanine coupling, whereas CHI animals showed little or no detectable association. Alongside the disrupted 3PG-to-pyruvate and pyruvate-to-lactate relationships, this pattern is consistent with impaired carbon metabolism at the pyruvate branchpoint in chronic KI CHI hippocampus, with a similar pyruvate-to-alanine pattern in cortex.

**Figure 5:**
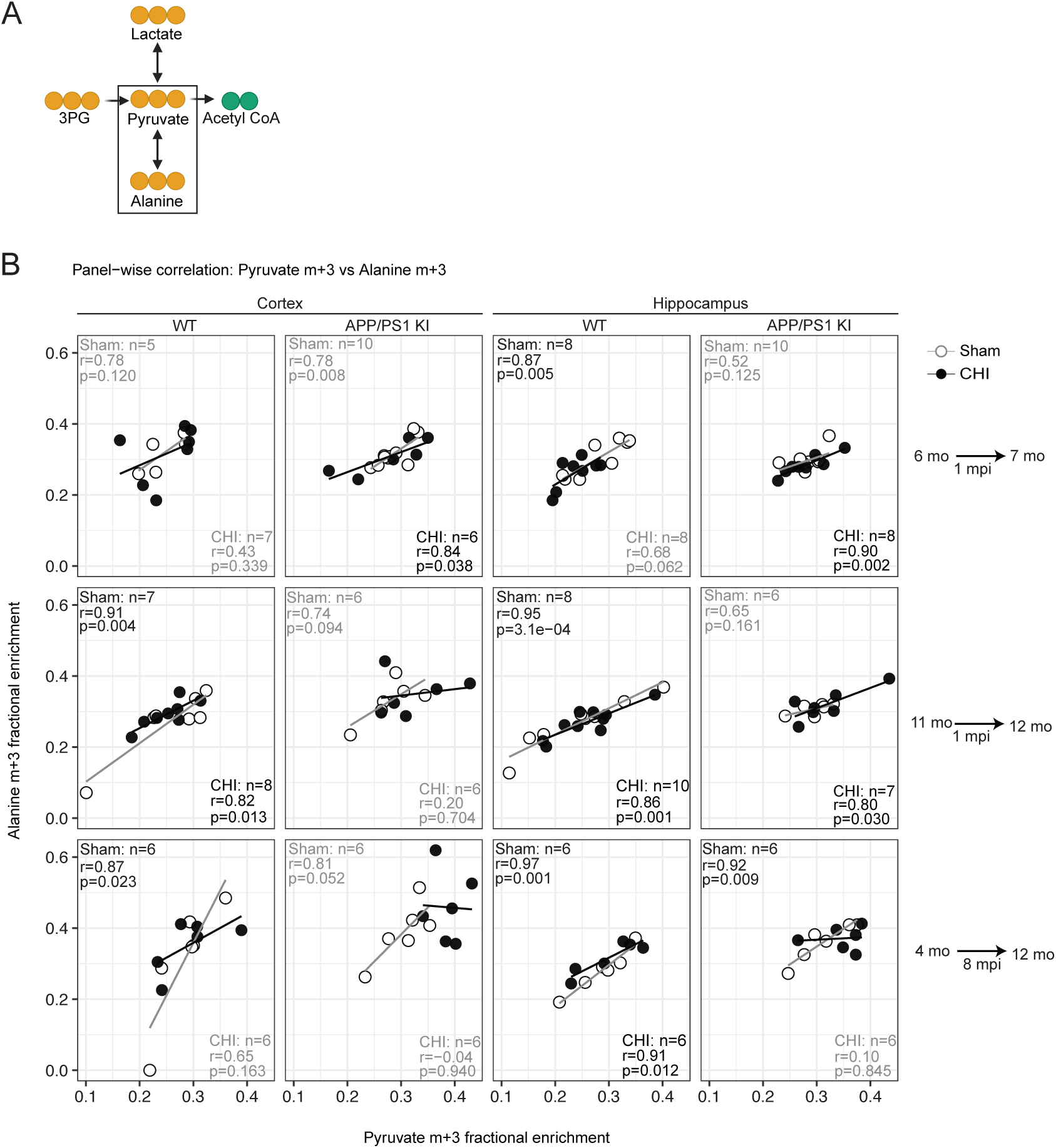
Weakened pyruvate-to-alanine coupling in KI hippocampus after CHI. **(A)** Schematic showing alanine as a transamination-linked fate of pyruvate at the pyruvate branchpoint. (**B**) Panel-wise correlations between pyruvate m+3 and alanine m+3 fractional enrichment in cortex and hippocampus from WT and KI mice under sham or CHI conditions across three cohorts: mature-adult 1-month post-injury (7 months at endpoint), middle-aged 1-month post-injury (12 months at endpoint), and chronic 8-months post-injury (mature-adult-injured, 12 months at endpoint). Sham-versus-CHI differences in correlation strength were assessed by Fisher’s r-to-z comparison, and differences in regression slope were assessed using a slope-interaction linear model (y ∼ x * surgery). Abbreviations: mpi, month post-injury; mo, months.

To determine whether the chronic abnormality extended beyond the pyruvate branchpoint into downstream oxidative metabolism, we next examined precursor-product pairs spanning sequential steps in first-turn TCA-cycle labeling. Pyruvate m+3 versus citrate m+2 reflects entry of pyruvate-derived carbon into the cycle through acetyl-CoA condensation with oxaloacetate (Fig. S2). Citrate m+2 to glutamate m+2 follows that carbon onward through α-ketoglutarate and its amino-acid pool (Fig. S3). The glutamate-to-GABA relationship (Fig. S4) tests whether downstream neurotransmitter coupling is preserved, and the malate-to-aspartate relationship (Fig. S5) reports coupling between later TCA intermediates and oxaloacetate-linked transamination. Across these analyses, the most consistent loss of coupling was at pyruvate-to-citrate, with positive associations concentrated in WT hippocampus and weak or absent coupling across most cortical panels (Fig. S2). Citrate-to-glutamate coupling was generally maintained, although in chronic KI hippocampus it reversed to a negative correlation after CHI at a trend level (Fig. S3). By contrast, glutamate-to-GABA and malate-to-aspartate coupling remained strongly positive across nearly all groups (Figs. S4–S5). Taken together, the chronic KI CHI pattern was specific to 3PG-to-pyruvate loss, pyruvate-to-lactate loss, and pyruvate-to-alanine weakening in chronic KI hippocampus.

### 3.6 Histological and biochemical assessment of amyloid-associated pathology

To determine whether CHI increased the amyloid burden, we performed multiplex immunofluorescence with Amylo-Glo, IBA1, GFAP and CD45 (Fig. 6A-B, single-channel images in Fig. S6). We measured Amylo-Glo-positive plaque area in neocortex and hippocampus of APP/PS1 KI mice, with no detectable plaques in wild type mice. There was no increase in Aβ plaque area in either region following injury (Fig. 6C-D). Since PBS- and formic acid-soluble fractions report different Aβ pools, we extracted cortical tissue sequentially to separate them. The PBS fraction recovers monomeric and oligomeric Aβ that remains soluble in aqueous buffer, whereas the subsequent formic acid fraction solubilizes fibrillar, deposited Aβ. Protein-normalized Aβ42 did not differ significantly between KI CHI and KI sham mice in any cohort after BH correction (Fig. 6E), including the only nominal difference, lower PBS Aβ42/protein in KI CHI mice at 1 month post-injury (7 months old; p=0.036, q=0.108).

**Figure 6:**
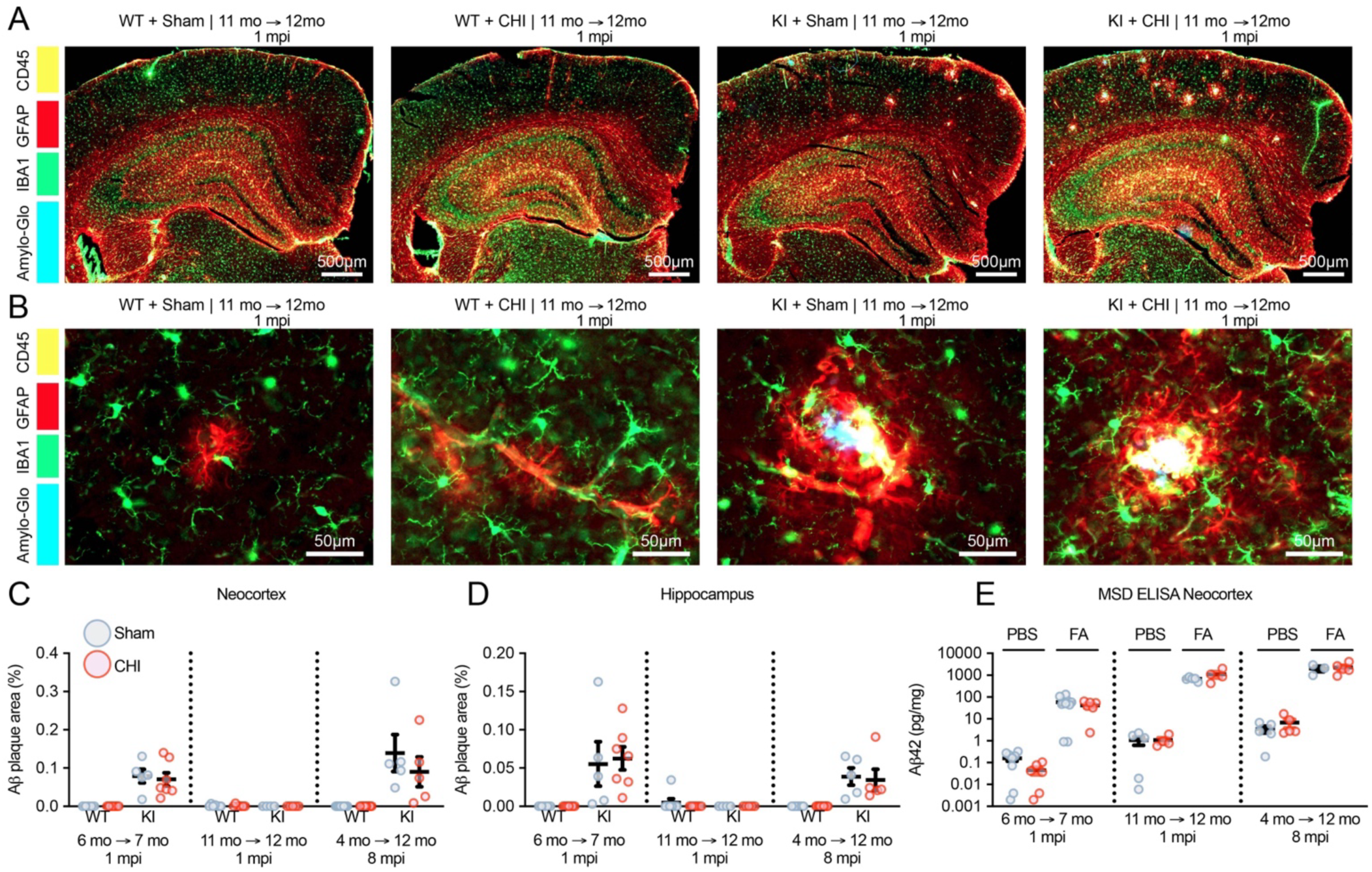
Representative multiplex immunofluorescence, regional Aβ plaque burden, and cortical Aβ42 measurements after CHI. (**A-B**) Representative multiplex immunofluorescence from WT sham, WT CHI, KI sham and KI CHI mice. Aβ signal is shown in blue, GFAP in red, IBA1 in green and CD45 in yellow. Images were acquired at 40x magnification; scale bar, 10 μm. (**C**) Aβ-positive plaque area as a percentage of the analyzed neocortical region across the mature-adult 1-mo post-injury (7 months at endpoint), middle-aged 1-mo post-injury (12 months at endpoint), and chronic 8-mo post-injury (12 months at endpoint) cohorts. (**D**) Corresponding Aβ-positive plaque area in hippocampus. (**E**) PBS-soluble and formic-acid-soluble cortical Aβ42 measured by singleplex MSD and normalized to PBS-fraction protein concentration, reported as pg Aβ42/mg protein. Sham mice are shown as gray circles and CHI mice as red open circles; horizontal bars indicate group means ± SEM. Each point represents one biologically independent mouse; sample sizes vary across MSD fractions because of prespecified assay-quality exclusions. Statistical results are reported in Supplemental table 2. Abbreviations: mpi, month post-injury; mo, months.

To determine whether CHI altered overall glial response, we quantified marker-positive area across whole-region annotations in neocortex and hippocampus (Fig. 7A-F). In the chronic 8-month post-injury cohort (12 months at endpoint), neocortical GFAP area showed an injury effect (p=0.032; Fig. 7A). The direct KI CHI-versus-KI sham comparison was not significant by Mann-Whitney testing (p=0.222). In the middle-aged 1-month post-injury cohort (12 months old), hippocampal GFAP area showed genotype x injury interaction (p=0.042) and an injury effect (p=0.041), while hippocampal CD45 area also showed an injury effect (p=0.0064). The direct KI CHI-versus-KI sham comparison for hippocampal CD45 was not significant (p=0.6169).

**Figure 7:**
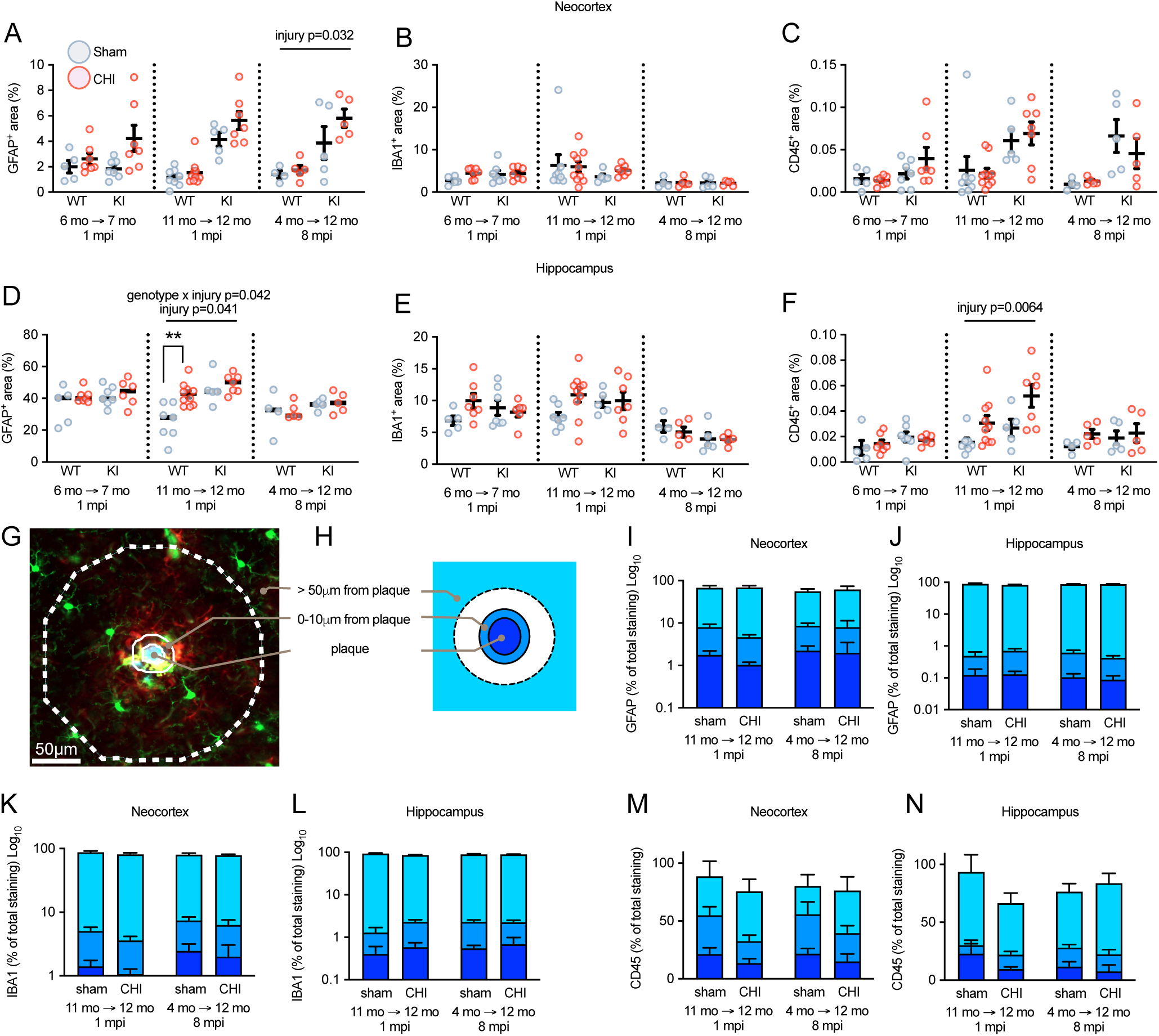
Region-wide glial-marker area and plaque-compartment distribution after CHI. **(A-C)** Whole-region marker-positive area in neocortex for (**A**) GFAP, (**B**) IBA1, (**C**) CD45, each shown for all three cohorts, comprising mature-adult 1 mpi (7 months at endpoint), middle-aged 1 mpi (12 months at endpoint), and chronic 8 mpi (12 months at endpoint). (**D-F**) Corresponding measurements in hippocampus for (**D**) GFAP, (**E**) IBA1, and (**F**) CD45. Each point is one animal, averaged across two sections. Sham mice are shown as gray open circles and CHI mice as red open circles, with horizontal black bars indicating group mean ± SEM. (**G**) Representative neocortex multiplex immunofluorescence image of a single plaque with concentric partitioners overlaid from chronic 8pi KI +CHI. Channel assignments are as in Supplemental Figure 6. Magnification 50x, scale bar 50 μm. (**H**) Schematic of the plaque-associated compartments, comprising the plaque core, 0-10 μm from the plaque perimeter, and tissue more than 50 μm from the plaque. (**I-N**) Distribution of marker-positive staining across plaque compartments in KI mice, for GFAP in (**I**) neocortex and (**J**) hippocampus, and CD45 in (**M**) neocortex and (**N**) hippocampus. Values are the percentage of each animal’s marker-positive area within each brain region, with error bars indicating mean ± SEM. GFAP and IBA1 are plotted on log_10_ y-axis. CD45 is plotted on a linear percentage scale. Abbreviations: mpi, month post-injury; mo, months.

Microglia cluster around fibrillar Aβ deposits in AD brain (Condello et al., 2015). To determine whether CHI altered the glial response to amyloid pathology, we partitioned marker-positive staining in KI mice into the plaque core, 0-10 μm peri-plaque annulus, and tissue more than 50 μm from the plaque (Fig. 7G-H). In both neocortex and hippocampus, the large majority of GFAP-, IBA1, and CD45-positive staining was located in the distal (50 μm) compartment, consistent with the small fraction of tissue area occupied by plaques and their immediate surround. This distribution was smaller between KI sham and KI CHI mice in the middle-aged 1-month and chronic 8-month post-injury cohorts (Fig.7I-N). CHI therefore did not detectably shift glial marker burden toward or away from plaques at either post-injury interval.

## 4. Discussion

A central finding of this study is that a CHI in the APP/PS1 KI model produced a delayed shift in cerebral glucose-carbon handling. No broad labeling phenotype was evident 1 month post-injury, whereas clear regional changes were detected at 8 months. Specifically, KI cortex showed elevated pyruvate m+3 and lactate m+3, whereas KI hippocampus showed increased labeling from glucose m+6 through DHAP m+3 and 3PG m+3 to lactate m+3. First-turn TCA-linked labeling was not broadly reduced, but specifically 3PG-to-pyruvate and pyruvate-to-lactate coupling were lost in chronic KI CHI hippocampus. Thus, the chronic phenotype was centered on glycolytic and pyruvate/lactate nodes rather than a generalized loss of first-turn glucose-derived labeling.

Mitochondrial bioenergetics and first-turn glucose-carbon labeling capture different aspects of post-injury metabolism. In our prior study using the same model, the strongest genotype-by-injury effects on mitochondrial function occurred at 1 month after CHI, particularly in the hippocampus (Moallem et al., 2026). However, no corresponding broad change in first-turn labeling was detected at 1-month. Altered glucose-derived labeling emerged at 8 months, when mitochondrial function was no longer impaired in our prior study. This temporal dissociation suggests that early mitochondrial dysfunction may contribute to a delayed metabolic response rather than an immediate altering of first-turn glucose-derived labeling. An intervention that selectively prevents the early bioenergetic deficit could test whether it contributes to the chronic labeling phenotype.

Pyruvate is the main oxidative gate linking glycolysis to mitochondrial oxidation, with PDH controlling its conversion to acetyl-CoA. Studies in TBI and AD-related brain tissue implicate PDH dysfunction, while isotope tracing after cortical injury has shown altered distribution of glucose-derived carbon through PDH- and pyruvate carboxylase-linked pathways despite continued oxidative labeling (Bartnik, Hovda, et al., 2007; Bubber et al., 2005; Hoshi et al., 1996; Lazzarino et al., 2019; Sharma et al., 2009; Sheu et al., 1985; Xing et al., 2009). The chronic cortical increase in pyruvate m+3 and lactate m+3, without a broad reduction in TCA-linked labeling, is consistent with a partial constraint or redistribution at the pyruvate oxidative gate rather than complete failure of glucose-derived carbon entry into the TCA cycle.

Lactate accumulation and elevated lactate/pyruvate ratios are well described after TBI, but these findings can reflect increased glycolysis, altered redox balance, or impaired lactate utilization or clearance and therefore do not identify a single mechanism (Brooks & Martin, 2014; Carpenter et al., 2015; Jalloh et al., 2018; Lekomtseva, 2020; Patet et al., 2016; Sahuquillo et al., 2014). In chronic KI hippocampus, the loss of pyruvate-to-lactate and 3PG-to-pyruvate coupling indicates that the phenotype extended beyond increased lactate labeling to disruption of normal relationships among glycolytic precursors, pyruvate, and lactate.

The chronic KI CHI hippocampus phenotype also extended upstream of pyruvate, with increased glucose m+6, DHAP m+3, and 3PG m+3 but no significant increase in pyruvate m+3. This pattern is consistent with altered handling of glucose-derived carbon before the pyruvate branchpoint and it could involve changes in glycolytic processing or PPP activity, as previously reported after TBI (Bartnik, Lee, et al., 2007; Carpenter et al., 2014; Jalloh et al., 2015). However, [U ^13^-C]-glucose fractional enrichment cannot distinguish glycolytic recycling from PPP shunting or established that flux through either pathway was increased.

The chronic KI CHI pattern differs from the broader reduction in downstream glucose-derived labeling reported in APP/PS1 transgenic at 10 months, in which [1-^13^C]-glucose produced lower ^13^C enrichment across TCA-cycle intermediates and neurotransmitter-linked metabolites (Zhou et al., 2018), and from the age-dependent shift between hyper- and hypometabolic TCA-cycle labelling in 3xTg-AD mice (Sancheti, Kanamori, et al., 2014; Sancheti, Patil, et al., 2014). The human brain oxidizes lactate to glutamine via the TCA cycle (Gallagher et al., 2009). In injured brain, however, extracellular pyruvate and lactate can rise while labelled lactate is recovered with no detectable ^13^C enrichment of TCA-cycle-derived glutamine (Stovell et al., 2023). Upstream and oxidative labeling can therefore move independently. Non-tracer ^1^H-NMR metabolomics in APP23/PS45 mice subjected to TBI likewise identified changes in glycolytic and TCA-metabolites, though injury also accelerated β-secretase processing and Aβ deposition, which we did not observe (Lou et al., 2018).

The chronic metabolic changes were not accompanied by a generalized increase in Aβ42 or glial-marker staining. Regional glial responses were detected, but were not uniform across brain regions or markers, and no direct KI CHI-versus-KI sham comparison reached significance. Protein normalized cortical Aβ42 did not differ between KI CHI and KI sham mice at any post-injury endpoint after correction for multiple comparisons. Altered handling of glucose-derived carbon therefore does not appear to be a simple downstream consequence of amyloid accumulation or of a generalized increase in glial marker burden and may instead reflect a persistent injury effect on metabolic regulation in the KI mice. Consistent with the upstream component of this pattern, [U-^13^C]-glucose labelling of lactate and alanine is elevated in APPswe/PSEN1dE9 cortical slices at 3 months, before plaque deposition, although TCA-cycle labelling was concurrently reduced (Andersen et al., 2017) and microglial metabolic programs shift in APP/PS1 mice by 2 months, before morphological activation or plaque formation is apparent (Qiu et al., 2021). Because glial marker area is not a measure of glial metabolic state, metabolically reprogrammed glia could still have contributed to the observed labeling changes (Baik et al., 2019; Pan et al., 2022).

Several limitations define the scope of these findings. Fractional enrichment and precursor-product coupling do not measure absolute fluxes or enzyme activity. Direct PDH and LDH activity assays, monocarboxylate-transporter measurements, and kinetic tracer studies are therefore needed to test the proposed branchpoint mechanism. The 30-minute sampling point captures first-turn labeling but not later TCA-cycle turns or slower exchange pools; serial sampling would define their time courses. Although two cohorts shared a 12-month endpoint, post-injury interval cannot be fully separated from the time available for aging and amyloid-related pathology to progress. An expanded design spanning multiple ages at injury and post-injury intervals would better distinguish these influences. Bulk-tissue measurements also obscure neuronal, astrocytic, and vascular contributions, which could be addressed through cell-resolved isotope tracing. Finally, because only male mice were available for this study, the chronic phenotype must also be tested in females.

## 5. Conclusions

The chronic metabolic consequence of CHI in the KI model was a delayed and selective disruption in glucose-carbon routing rather than a global loss of first-turn oxidative labeling. This shift occurred without a corresponding increase in amyloid burden or a generalized glial response, despite clear regional changes in glucose-derived labeling. By identifying enrichment patterns consistent with altered carbon fate at the pyruvate branchpoint, the present study identifies a candidate site of vulnerability for future mechanistic studies and metabolic interventions after injury in AD-relevant brain.

## Author Contributions

Elika Z. Moallem contributed to writing the original draft, visualization, data analysis, and investigation. Ronald C. Bruntz contributed to methodology, data curation, investigation, and writing – review and editing. Samir P. Patel contributed to data analysis, interpretation, and writing - review and editing. MaKayla F Cox participated in investigation and writing – review and editing. Leke Bytyqi participated in investigation and writing – review and editing. Taylor K. Roberts participated in investigation and writing – review and editing. Teresa Macheda contributed to writing - review and editing and project administration. Kelly N. Roberts participated in investigation and writing - review and editing. Emma K. Higgins participated in investigation and writing – review and editing. Ramon C. Sun contributed to interpretation, funding acquisition, and writing - review and editing. Matthew S. Gentry contributed to interpretation, funding acquisition, and writing - review and editing. Patrick G. Sullivan contributed to data analysis, interpretation, funding acquisition, and writing - review and editing. Lance A. Johnson contributed to data analysis, interpretation, funding acquisition, and writing - review and editing. Adam D. Bachstetter was responsible for conceptualization, funding acquisition, project administration, writing the original draft, and data interpretation.

## Data Availability

Data used in this study are available through the Open Data Commons for Traumatic Brain Injury.

## Declaration of Generative AI Usage in Scientific Writing

During the preparation of this work, the author(s) used ChatGPT and Claude to assist in evaluating the original text for clarity, completeness, and style. These tools provided editorial suggestions and recommendations; however, the author(s) reviewed and edited all content as necessary. The author(s) take full responsibility for the final content of the published article.

## Abbreviations

3PG: 3-phosphoglycerate;
AD: Alzheimer’s disease;
ANOVA: analysis of variance;
APP/PS1: amyloid precursor protein/presenilin 1;
ARRIVE: Animal Research: Reporting of In Vivo Experiments;
CHI: closed head injury;
DHAP: dihydroxyacetone phosphate;
DExSI: Data Extraction for Stable Isotope-labelled metabolites;
DLST: dihydrolipoamide S-succinyltransferase;
EI: electron ionization;
FAD: familial Alzheimer’s disease;
GABA: γ-aminobutyric acid;
GC-MS: gas chromatography-mass spectrometry;
KI: knock-in;
LDH: lactate dehydrogenase;
MANOVA: multivariate analysis of variance;
mo: months;
mpi: month post-injury;
MSTFA: N-methyl-N-trimethylsilyl-trifluoroacetamide;
PDH: pyruvate dehydrogenase;
PPP: pentose phosphate pathway;
RRID: Research Resource Identifier;
SD: standard deviation;
SEM: standard error of the mean;
TBI: traumatic brain injury;
TCA: tricarboxylic acid cycle;
[U-^13^C]-glucose: uniformly ^13^C-labeled glucose;
WT: wild-type.

