## Supplemental Figures for "Closed-head injury produces a delayed, region-specific defect in glucose-carbon routing at the pyruvate branchpoint in APP/PS1 KI mice"

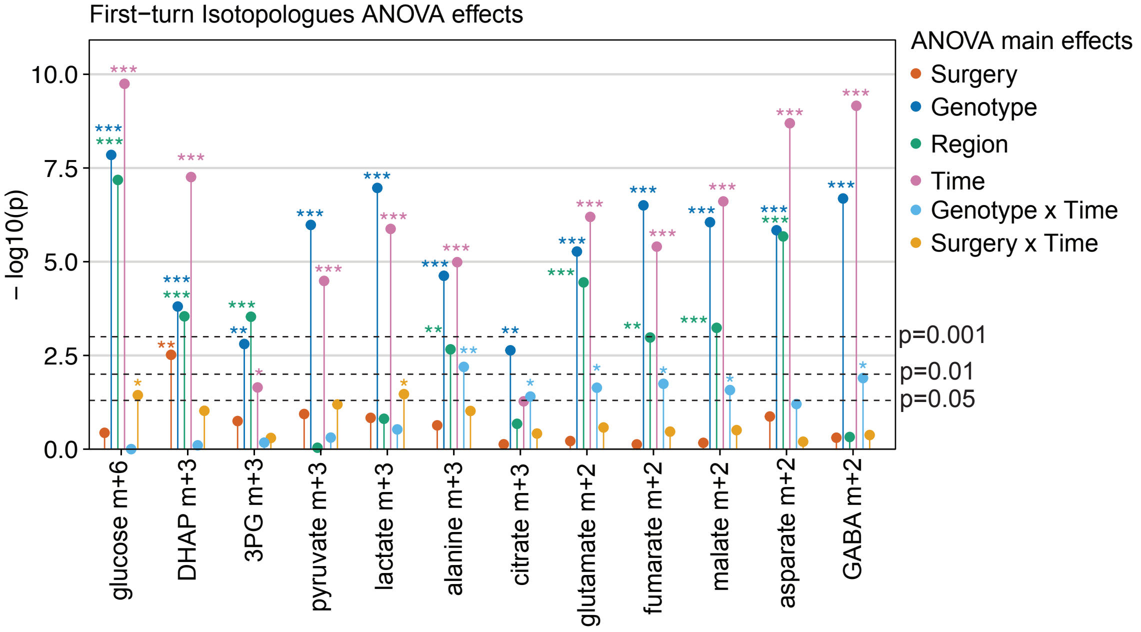


**Supplemental Figure S1: First-turn isotopologue effects identified by factorial ANOVA.** Lollipop showing -log10 (nominal p) values from univariate Type III ANOVAs for each first-turn isotopologue (x-axis). Colors identify the ANOVA terms shown: the main effects of surgery, genotype, brain region, and post-injury time, together with interaction terms retained for display when at least one isotopologue had a nominal $p<0.05$. Horizontal dashed lines indicate nominal p=0.05, 0.01, and 0.001 thresholds. Values are unadjusted p-values and indicate statistical evidence rather than effect size or direction.

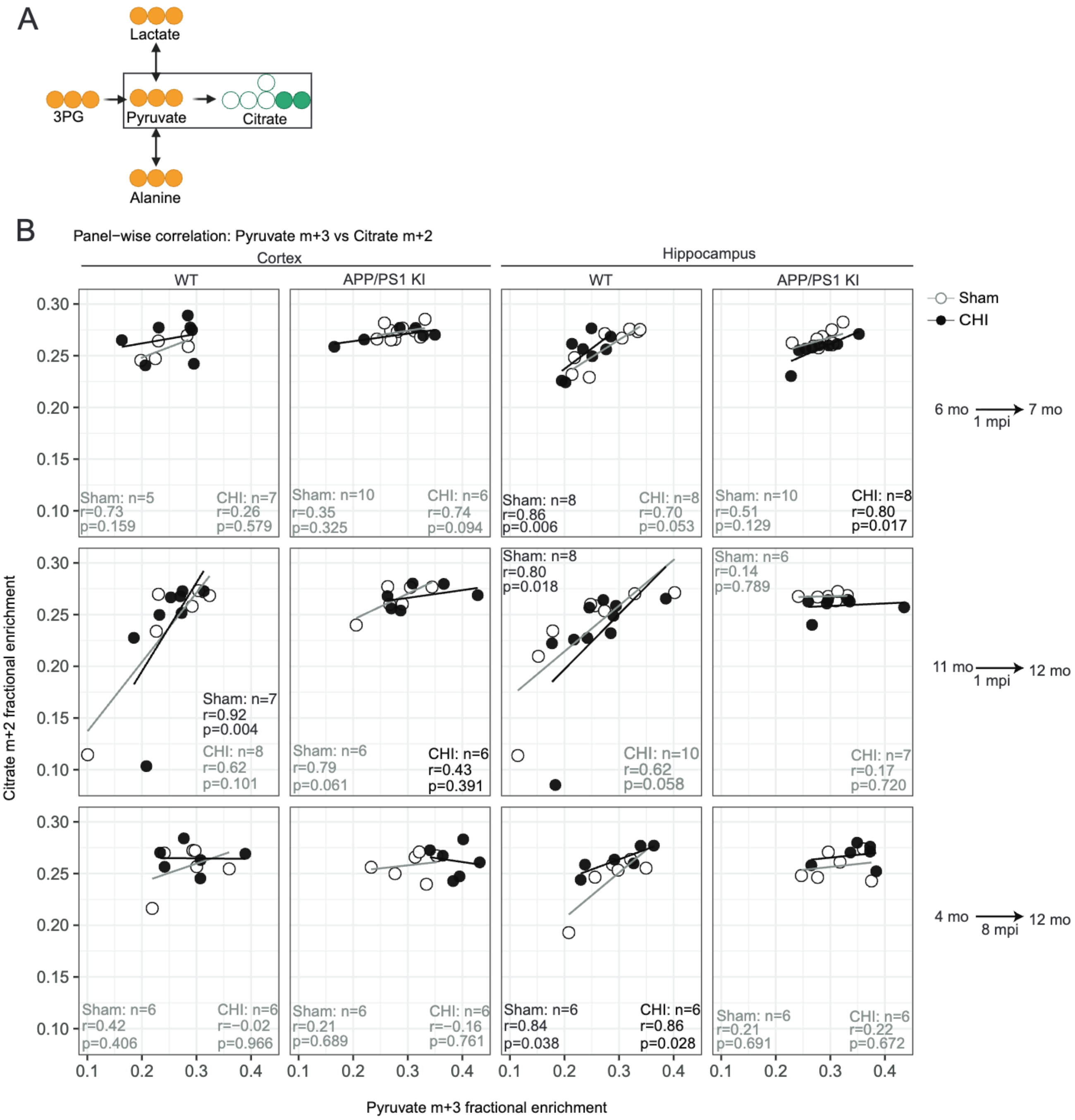


**Supplemental Figure S2: Pyruvate-to-citrate coupling across genotype, brain region, and post-injury cohort.** (**A**) Schematic showing citrate as the first-turn TCA-cycle product downstream of pyruvate entry through acetyl-CoA. (**B**) Panel-wise correlations between pyruvate m+3 and citrate m+2 fractional enrichment in cortex and hippocampus from WT and KI mice under sham or CHI conditions across three cohorts: mature-adult 1-month post-injury (7 months at endpoint), middle-aged 1-month post-injury (12 months at endpoint), and chronic 8-month post-injury (mature-adult-injured, 12 month at endpoint). Sham-versus-CHI differences in correlation strength were assessed by Fisher's r-to-z comparison, and differences in regression slope were assessed using a slope-interaction linear model (y ~ x * surgery). Abbreviations: mpi, month post-injury; mo, months.

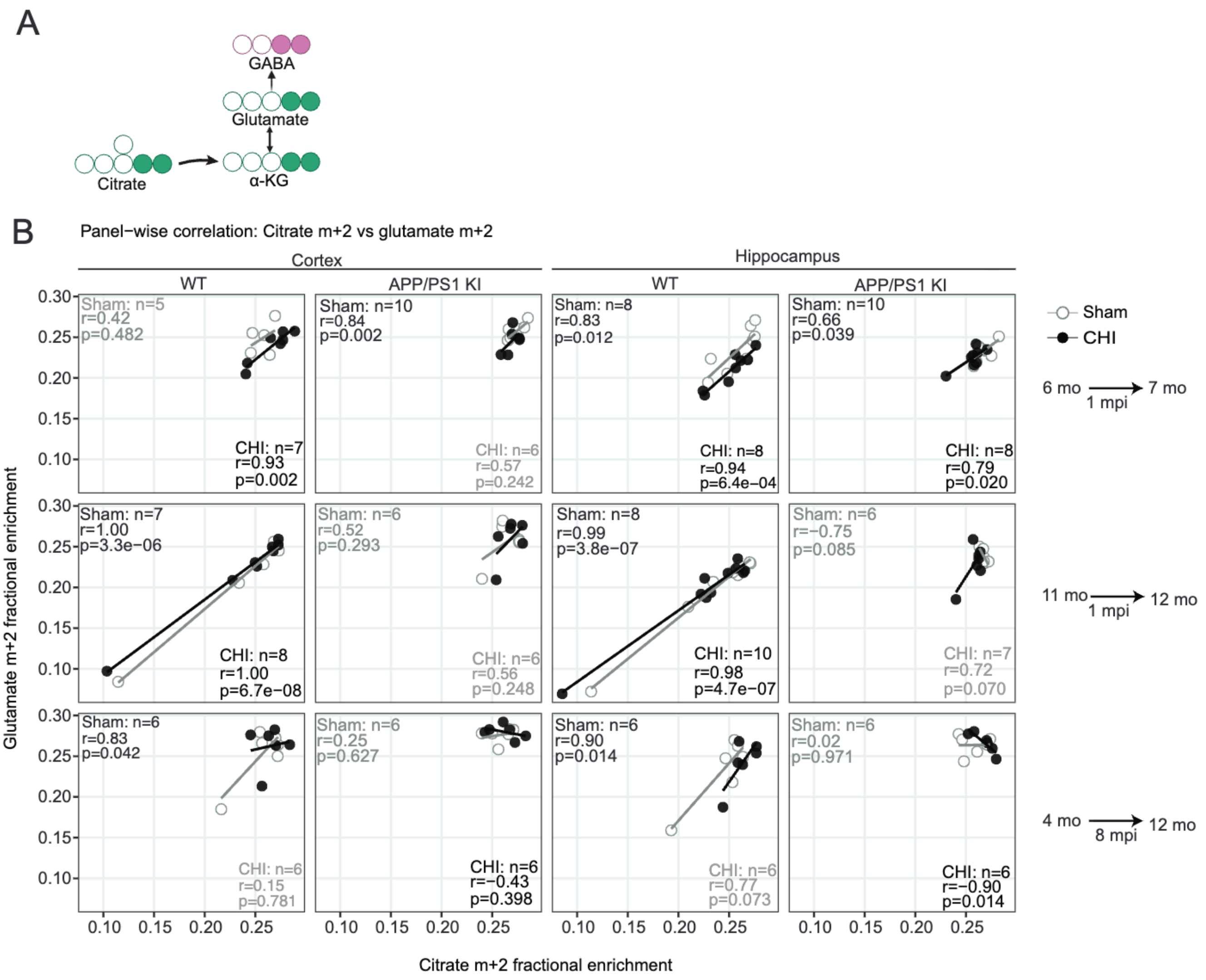


**Supplemental Figure S3: Citrate-to-glutamate coupling across genotype, brain region, and post-injury cohort.** (**A**) Schematic showing glutamate as a downstream amino acid pool linked to first-turn TCA-cycle labeling through citrate and α-ketoglutarate. (**B**) Panel-wise correlations between citrate m+2 and glutamate m+2 fractional enrichment in cortex and hippocampus from WT and KI mice under sham or CHI conditions across three cohorts: mature-adult 1-month post-injury (7 months at endpoint), middle-aged 1-month post-injury (12 months at endpoint), and chronic 8-month post-injury (mature-adult-injured, 12 months at endpoint). Sham-versus-CHI differences in correlation strength were assessed by Fisher’s r-to-z comparison, and differences in regression slope were assessed using a slope-interaction linear model (y ~ x * surgery). Abbreviations: mpi, month post-injury; mo, months.

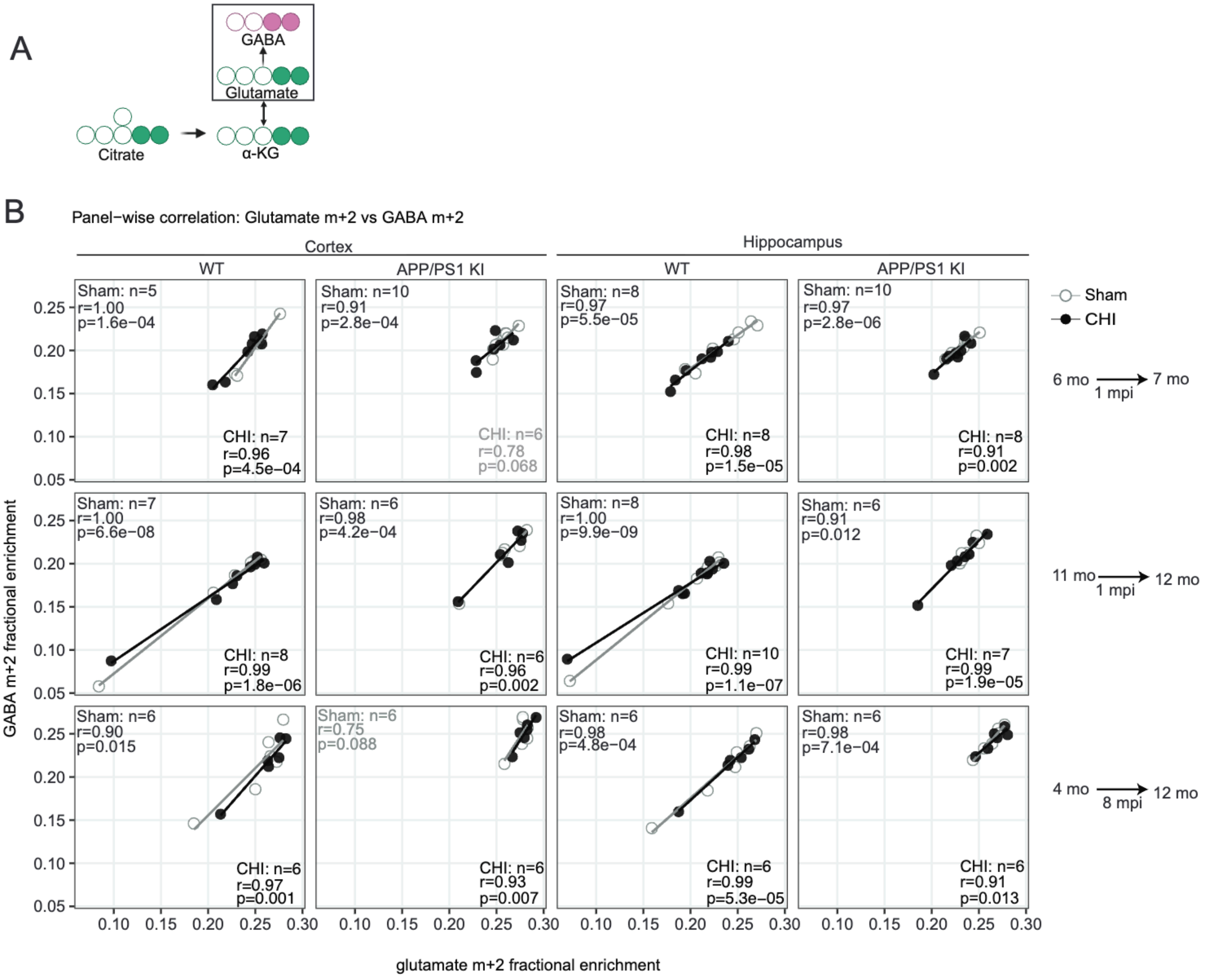


**Supplemental Figure S4: Glutamate-to-GABA coupling across genotype, brain region, and post-injury cohort.** (**A**) Schematic showing GABA as a downstream neurotransmitter-linked fate of glutamate. (**B**) Panel-wise correlations between glutamate m+2 and GABA m+2 fractional enrichment in cortex and hippocampus from WT and KI mice under sham or CHI conditions across three cohorts: mature-adult 1-month post-injury (7 month at endpoint), middle-aged 1-month post-injury (12 months at endpoint), and chronic 8-month post-injury (mature-adult-injured, 12 months at endpoint). Sham-versus-CHI differences in correlation strength were assessed by Fisher's r-to-z comparison, and differences in regression slope were assessed using a slope-interaction linear model (y ~ x * surgery). Abbreviations: mpi, month post-injury; mo, months.

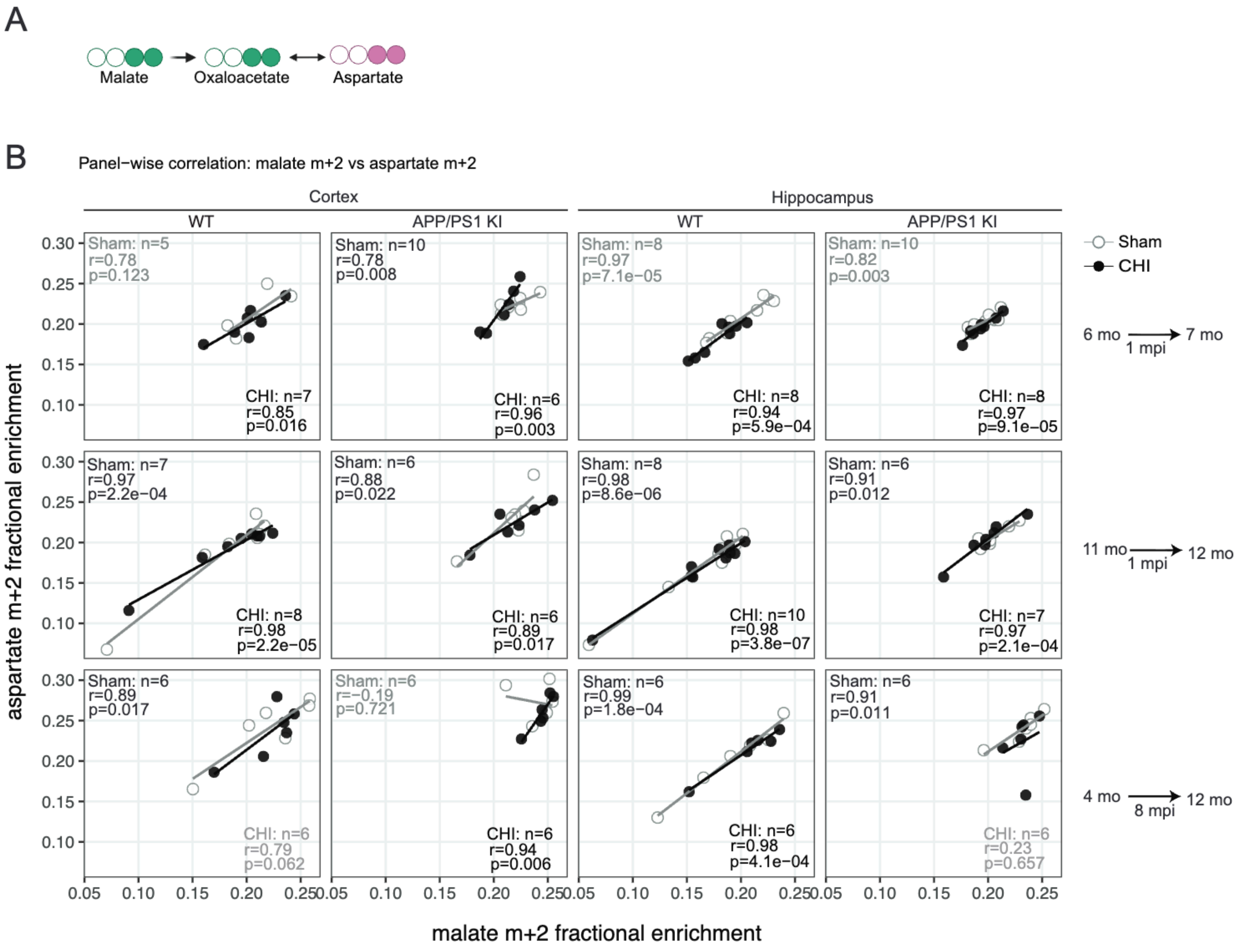


**Supplemental Figure S5: Malate-to-aspartate coupling across genotype, brain region, and post-injury cohort.** (A) Schematic showing aspartate as a downstream transamination-linked product of oxaloacetate derived from malate. (B) Panel-wise correlations between malate m+2 and aspartate m+2 fractional enrichment in cortex and hippocampus from WT and KI mice under sham or CHI conditions across three cohorts: mature-adult 1-month post-injury (7 months at endpoint), middle-aged 1-month post-injury (12 months at endpoint), and chronic 8-month post-injury (mature-adult-injured, 12 months at endpoint). Sham-versus-CHI differences in correlation strength were assessed by Fisher's r-to-z comparison, and differences in regression slope were assessed using a slope-interaction linear model (y ~ x * surgery). Abbreviations: mpi, month post-injury; mo, months.

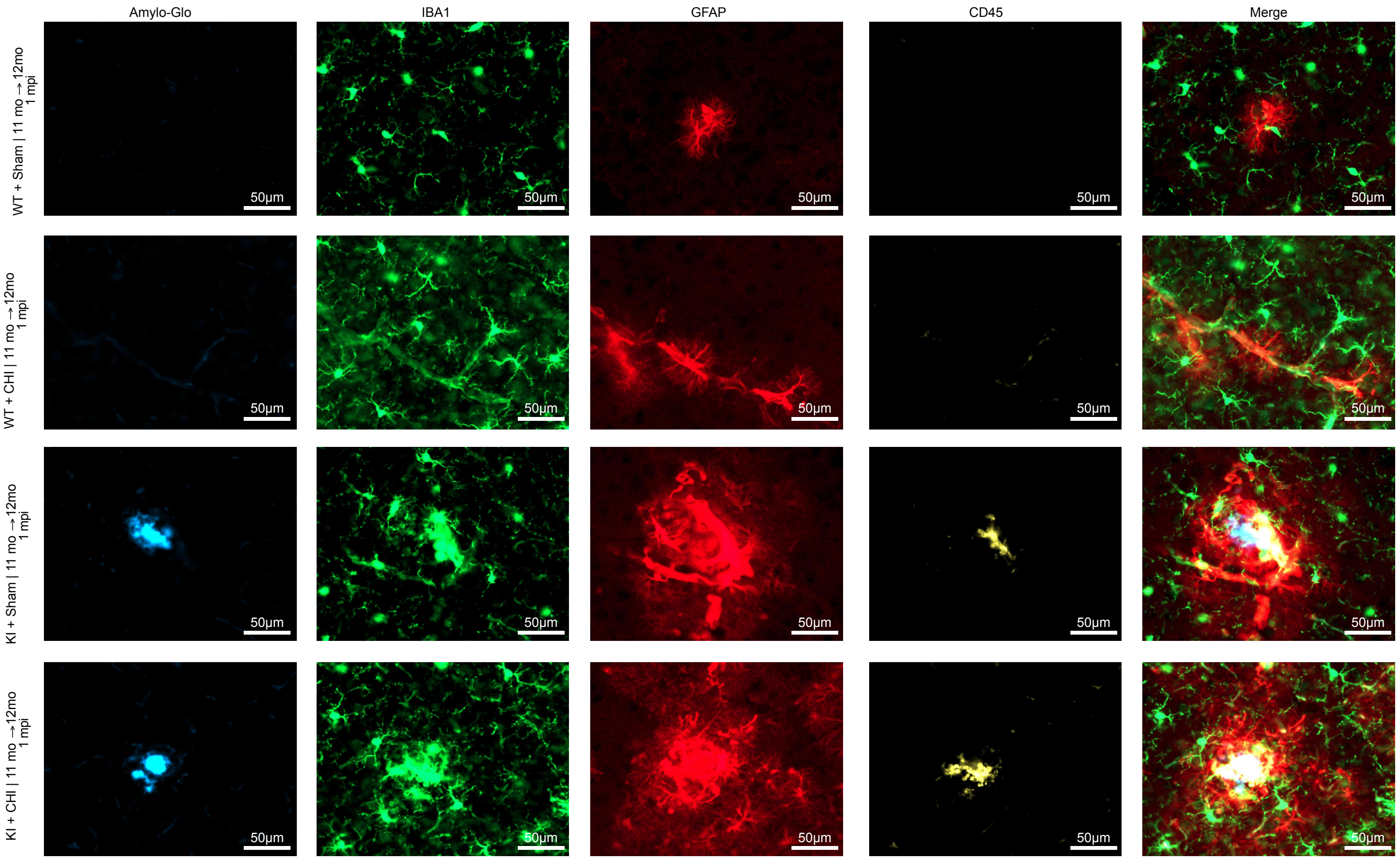


**Supplemental Figure S6: Single-channel and merged multiplex immunofluorescence for amyloid plaques, microglia, astrocytes, and CD45 in WT and APP/PS1 KI mice 1 month after closed-head injury.** Free-floating coronal sections (30 µm) from WT sham, WT CHI, KI sham, KI CHI mice of the middle-aged 1-month post-injury cohort (12 months at endpoint) were stained with Amylo-Glo (blue), IBA1 (green), GFAP (red), and CD45 (yellow), and imaged on a Zeiss AxioScan Z.1. Each row shows each experimental group; columns show the individual channels followed by the four-channel merge. Amylo-Glo positive deposits are present only in KI mice; the WT panels show the corresponding fields in the absence of amyloid plaques. Images were acquired at 40x magnification from cortex; scale bar 10 µm. Images are shown for illustration. All quantification was performed on the full outlined section as described in Methods. Abbreviations: mpi, month post-injury; mo, months.
